# Conditional antagonism in co-cultures of *Pseudomonas aeruginosa* and *Candida albicans*: an intersection of ethanol and phosphate signaling distilled from dual-seq transcriptomics

**DOI:** 10.1101/2020.04.20.050765

**Authors:** Georgia Doing, Katja Koeppen, Patricia Occipinti, Deborah A. Hogan

## Abstract

*Pseudomonas aeruginosa* and *Candida albicans* are opportunistic pathogens whose interactions involve the secreted products ethanol and phenazines. Here we describe the focal role of ethanol in mixed-species co-cultures by dual RNA-seq analyses. *P. aeruginosa* and *C. albicans* transcriptomes were assessed after growth in mono-culture or co-culture with either ethanol-producing *C. albicans* or a *C. albicans* mutant lacking the primary ethanol dehydrogenase, Adh1. Analyses using KEGG-pathways and the previously published eADAGE method revealed several *P. aeruginosa* responses to *C. albicans*-produced ethanol including the induction of a non-canonical low phosphate response mediated by PhoB. *C. albicans* wild-type, but not *C. albicans adh1*Δ/Δ, induces *P. aeruginosa* production of 5-methyl-phenazine-1-carboxylic acid (5-MPCA), which forms a red derivative within fungal cells. We first demonstrate that PhoB is required for this interaction and that PhoB hyperactivity, via deletion of *pstB,* leads to increased production of 5-MPCA even when phosphate concentrations are high, but only in the presence of ethanol. Second, we show that ethanol is only sufficient to promote 5-MPCA production at permissive phosphate concentrations. The intersection of ethanol and phosphate in co-culture is mirrored in *C. albicans*; the *adh1*Δ/Δ mutant had increased expression of genes regulated by Pho4, the *C. albicans* transcription factor that responds to low phosphate which we confirmed by showing the *adh1*Δ/Δ strain had elevated Pho4-dependent phosphatase activity. The dual-dependence on ethanol and phosphate concentrations for anti-fungal production highlights how environmental factors modulate microbial interactions and dictate antagonisms such as those between *P. aeruginosa* and *C. albicans*.

**Author Summary:** *Pseudomonas aeruginosa* and *Candida albicans* are opportunistic pathogens that are frequently isolated from co-infections. Using a Dual-Seq approach in combination with genetics approaches, we found that ethanol produced by *C. albicans* stimulates the PhoB regulon in *P. aeruginosa* asynchronously with activation of the Pho4 regulon in *C. albicans.* In doing so, we demonstrate that eADAGE-based analysis can improve the understanding of the *P. aeruginosa* response to ethanol-producing *C. albicans* as measured by transcriptomics: we identify a subset of PhoB-regulated genes as differentially expressed in response to ethanol. We validate our result by showing that PhoB is necessary for multiple roles in co-culture including the competition for phosphate and the production of 5-methyl-phenazine-1-carboxylic acid, and that the *P. aeruginosa* response to *C. albicans*-produced ethanol depends on phosphate availability. The conditional stimulation of virulence production in response to sub-inhibitory concentrations of ethanol only under phosphate limitation highlights the importance of considering nutrient concentrations in the analysis of co-culture interactions.

## Introduction

*Pseudomonas aeruginosa* and *Candida albicans* are opportunistic pathogens that are frequently isolated from co-infections [1–11]. These pathogens affect each other’s behaviors through competition for nutrients [12–17], physical contact [3, 4, 7, 13, 14], diffusible signaling molecules [18–23] and antimicrobials [18, 21, 24–30]. Studies highlighting the dynamic interactions between *P. aeruginosa* and *C. albicans* have contributed to the growing understanding of how microbial interactions influence microbial physiology and behavior as well as microbiological and pathological outcomes.

Like many fermentative organisms, *C. albicans* produces ethanol. Ethanol is a biologically-active metabolite which, sub-inhibitory concentrations, modulates *P. aeruginosa* behavior in multiple ways: it induces activity of the sigma factor AlgU through ppGpp and DksA [31]; it promotes Pel matrix production through the Wsp system [29]; it decreases flagellar-mediated motility through a pathway implicated in cell-surface sensing [29, 32]; it affects pathways known to contribute to *P. aeruginosa* virulence [29, 33]; and it fuels fungal antagonism [29]. The broad effects of ethanol apply to many contexts and the response of *P. aeruginosa* to *C. albicans-*produced ethanol can serve as a model for how *P. aeruginosa* may respond to other fermentative fungi and bacteria. We seek to further understand this response and identify common themes which may be implicated in other microbial interactions.

To study the effects of ethanol in co-culture, we used *P. aeruginosa* anti-fungal production as a readout of the ethanol response. Previous work has shown that ethanol promotes the production and secretion of the phenazine 5-methyl-phenazine-carboxylic acid (5-MPCA) by *P. aeruginosa* and that, in turn, *P. aeruginosa* phenazines cause an increase in *C. albicans* fermentative metabolism and ethanol production [24–26]. *P. aeruginosa* does not normally secrete 5-MPCA in axenic cultures but in co-culture it secretes 5-MPCA through the MexGHI-OmpD efflux complex. Consequently, 5-MPCA enters *C. albicans* cells wherein it reacts with the amine group of arginine, disrupting protein function and forming a red pigment whose accumulation causes redox stress and eventually death of *C. albicans* [24, 27, 34]. While it is known that ethanol production by the fungus is necessary for *P. aeruginosa* 5-MPCA release, the mechanisms by which 5-MPCA production is regulated have not yet been described. The mechanisms of stimulation and ensuing consequences of *P. aeruginosa* 5-MPCA production and accumulation of the red 5-MPCA-derivative within *C. albicans* cells is a scopic case study for microbial interactions because it is an indicator of general antagonism.

Several studies have described the conditional production of antagonistic factors in response to nutrient availability such as phosphate or iron limitation [35–52]. This is often mediated transcriptionally, and such is the case for the low phosphate response which is mediated by the PhoR-PhoB two-component system wherein inorganic phosphate is sensed through the periplasmic domain of the phosphate transport complex, PstS, which is dependent on the ATPases PstA and PstB. The failure to bind phosphate in low phosphate environments causes the de-repression of the sensor kinase PhoR which phosphorylates the response regulator PhoB and initiates PhoB DNA-binding to the promoters of many genes.

This environmentally responsive regulation aids in the competition for essential nutrients. For example, the *P. aeruginosa* low-phosphate response includes the secretion of an arsenal of phosphatases, phospholipases and DNases that cleave phosphate from diverse macromolecules [40]. However, the secretion of these enzymes renders phosphate freely available to any nearby organism. Simultaneous production of antagonistic factors could aid *P. aeruginosa* in the competition for phosphate amongst other microbes. Indeed, in response to low phosphate and other complex stimuli, *P. aeruginosa* produces antagonistic factors like phenazines and phospholipases which play important roles in microbial interactions [39, 53, 54]. It has been reported that *P. aeruginosa* tailors its low phosphate response to secondary stimuli: in *P. aeruginosa* PhoB interacts with other regulators such as the transcription factor TctD [55] and the sigma factor VreI [37] to orchestrate the expression of its target genes. However, the mechanism by which PhoB exerts condition-specific control over its diverse regulon to manage antagonistic factors in microbial interactions, is not yet fully understood.

Co-culture transcriptomics data from single species RNA-Seq or dual RNA-Seq methods can be heterogeneous due to varying temporal and spatial relationships between organisms. Variability can make such data difficult to analyze with traditional statistical and pathway-based approaches. However, these data contain a wealth of information about nutrient competition, synergy and antagonism in microbial interactions. Therefore, it may be necessary to use techniques, such as recent machine learning based methods, that allow for the detection of subtle and novel transcriptional signals in order to render this important and complex data informative [53, 56–59].

Analysis of transcriptomic data from complex environments using curated pathways can be challenging if the conditions are not well represented by the data used for pathway definition. Furthermore, pathway definition relies on expert-contributed annotations, yet ∼38% (2,162 of 5,704) of genes for PAO1 reference strain (pseudomonas.com) lack description. Recent methods are using unsupervised machine learning to leverage large amounts of transcriptomic data and automatically identify sets of genes with correlated expression across large compendia of samples, agnostic of gene annotations and previously characterized pathways [56, 57, 59–62]. With over 2,000 transcriptional profiles of *P. aeruginosa* in the public sphere, such an approach has been successfully implemented to make expression-based gene sets which can be used as data-driven analytical tools that can bolster transcriptional analyses [53, 58, 63, 64]. In particular, the data-driven tool eADAGE has identified transcriptional signals that contain uncharacterized genes, manifest as small magnitude changes in expression, or are condition-specific yet biologically informative.

Here we demonstrate that *P. aeruginosa* and *C. albicans* undergo transcriptional changes in response to one another dependent on *C. albicans* ethanol production. Using eADAGE analysis we identify a group of PhoB-regulated genes as differentially expressed in response to ethanol and validate the result using genetic and biochemical assays that show PhoB is necessary for accumulation of the red 5-MPCA-derivative in *C. albicans* cells and that ethanol is sufficient to stimulate PhoB in mono-culture. We show that in co-culture PhoB regulation of phosphate scavenging and 5-MPCA production are independently necessary for *P. aeruginosa* fitness and antagonism against *C. albicans* respectively. Further, by examining *C. albicans* gene expression profiles matched to those of *P. aeruginosa* from the same co-cultures, we show that the *C. albicans* low phosphate response is inversely correlated to that of *P. aeruginosa* linking the coincidence of these stimuli in co-culture. By requiring dual-stimulation, In summary, we show that *P. aeruginosa* only produces antifungal 5-MPCA when the death of neighboring fungi would simultaneously remove a competitor and provide a source of the essential nutrient phosphate: in both the presence of fermenting *C. albicans* and phosphate limitation. We conclude that the enmity of *P. aeruginosa* – *C. albicans* interactions is conditional upon ethanol and phosphate concentrations.

## Results

### Ethanol is a defining factor in *P. aeruginosa – C. albicans* interactions stimulating antagonism wrought by transcriptional changes in both organisms

When grown on a lawn of *C. albicans*, *P. aeruginosa* produces the anti-fungal phenazine 5-MPCA which is taken up by *C. albicans* and modified within the fungal cells to form a red derivative [24, 27, 29] that can be seen first below and then as a halo surrounding *P. aeruginosa* colonies (**Fig. 1A**). The 5-MPCA precursor phenazine-1-carboxylic acid (PCA) can be synthesized via enzymes encoded in either of the two highly similar operons, *phzA1B1C1D1E1F1G1* (*phz1*) and *phzA2B2C2D2E2F2G2* (*phz2*) with different regulation; *phz1* contributes to phenazine production in liquid while *phz2* is responsible for phenazine production in colony biofilms [65]. Analysis of phenazine production in co-culture found that *phz1* was dispensable for the formation of red pigment, while *phz2* was required (**Fig. 1A**). Consistent with previous results, 5-MPCA-derived red pigment formation required *phzM* [27], *mexGHI-ompD* and *soxR* [24], and was over-abundant in upon deletion of *phzS,* which catalyzes the conversion of 5-MPCA into another phenazine, pyocyanin [66] (**Fig. S1A**). As we have previously reported and reproduced here, *P. aeruginosa* 5-MPCA production required *C. albicans* ethanol production as the *C. albicans adh1*Δ/Δ, which lacks the major ethanol dehydrogenase, did not elicit 5-MPCA-derived red pigment accumulation [29]. Chromosomal complementation of a single copy of *ADH1* in *C. albicans* restored *P. aeruginosa* 5-MPCA production (**Fig. 1A**).

**Figure 1.**
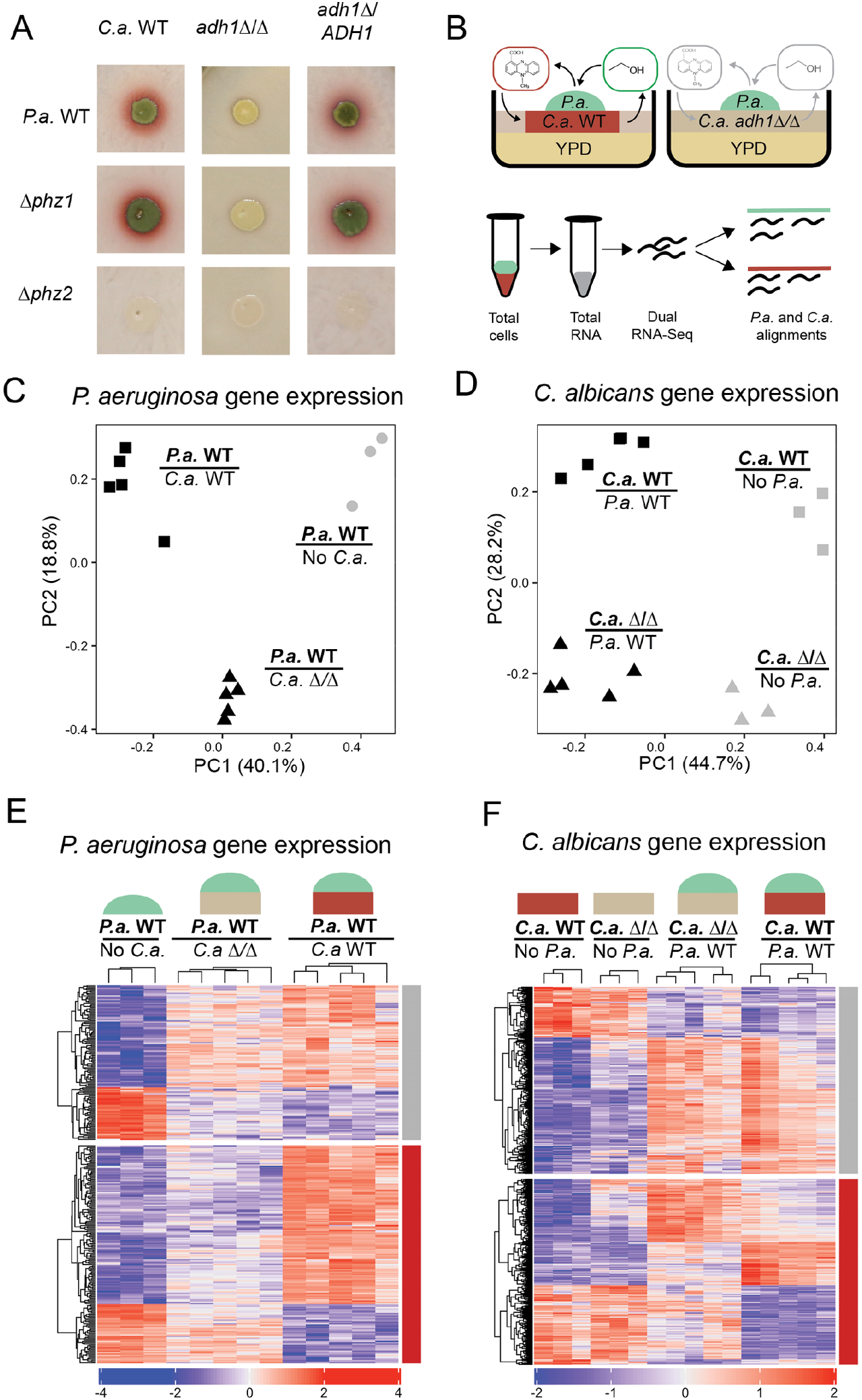
In co-culture of *C. albicans* (*C.a.*) and *P. aeruginosa* (*P.a*.), *C.a.*-produced ethanol stimulates *P.a.* to produce 5-MPCA and transcriptional responses ensue from both organisms. A) Co-cultures of *P.a*. wild type (WT) and mutants lacking phenazine biosynthesis operons (Δ*phz1* or Δ*phz2*) were inoculated onto 72 h-old lawns of *C.a.* wild type (WT), *C.a. adh1*Δ/Δ and *adh1*Δ/Δ reconstituted with *ADH1* (*adh1*Δ/*ADH1*). The red pigmentation indicates production of the phenazine 5-MPCA by *P.a..* B) Dual RNA-Seq allowed for parallel analyses of *P.a*. (green) and *C.a.* (red or pink) mRNA expression profiles from co-culture lawns to survey the effects of ethanol (green oval) and 5-MPCA (red oval) on gene expression. C) Principle component analysis (PCA) of TPM (transcripts per kilobase per million reads) from transcriptome profiles of *P.a.* grown alone (No *C.a.*), *P.a.* grown with *C.a.* WT, and *P.a.* grown with *C.a. adh1*Δ/Δ. D) PCA of gene expression profiles of *C.a.* WT and *C.a. adh1*Δ/Δ grown in mono-culture (No *P.a.*) or co-cultures with *P.a.* WT. E) The expression (z-score of TPM) of genes that differentiate *P.a.* in mono-culture from that grown in co-culture with *C.a.* WT (absolute value of log_2_fold-change (logFC) > 1 and false discovery rate (FDR) < 0.05); data for *P.a.* on *C.a*. *adh1*Δ/Δ are also shown. The red bar indicates genes that are significantly different between *C.a.* WT and *adh1*Δ/Δ (logFC >1, FDR < 0.05); the grey bar indicates genes that are not. F) Gene expression (z-score of TPM) of *C.a.* WT and *adh1*Δ/Δ grown in mono-culture or in co-culture with *P.a.*; genes that are significantly different between *C.a.* WT alone or *C.a.* WT with *P.a.* (logFC >1, FDR < 0.05) are shown for all four sample types. Genes that are also significantly different between *C.a.* WT and *adh1*Δ/Δ in the presence of *P.a*. (logFC >1, FDR < 0.05) are indicated by the red bar; the grey bar indicates genes that are not significantly different in this comparison.

To determine how *C. albicans* ethanol production influenced *P. aeruginosa* and how 5-MPCA-derived red pigment accumulation influenced *C. albicans*, we took a dual RNA-Seq approach in which we collected total RNA for simultaneous transcriptome-wide analyses of both organisms from 16 h co-cultures of *P. aeruginosa* with *C. albicans* WT, in which 5-MPCA-derivatives accumulated, and co-cultures of *P. aeruginosa* with *C. albicans adh1*Δ/Δ, in which 5-MPCA products were not observed (see **Fig. 1B** for experimental set up). Single-species *P. aeruginosa* and *C. albicans* colony biofilms grown on YPD medium were also analyzed at the same time point. Principle component analysis (PCA) of gene expression for each organism differentiated mono-culture and co-culture with the first component PC1, which accounted for 40.1% and 44.7% of total variance for *P. aeruginosa* and *C. albicans* respectively (**Fig. 1C,D**). The presence of *ADH1* in *C. albicans* constituted a defining feature of co-culture in PC2 for both organisms, which captured 18.8% and 28.2% of total variance for *P. aeruginosa* and *C. albicans* respectively (**Fig. 1C,D**). Comparison of *P. aeruginosa* gene expression on either *C. albicans* WT or *adh1*Δ/Δ to gene expression in mono-culture found 1,830 differentially expressed genes (DEGs) with an absolute log_2_fold-change (logFC) > 1 and a corrected p-value (FDR) < 0.05. Over half of the DEGs between *P. aeruginosa* in mono-culture and on *C. albicans* WT were also DEGs between *P. aeruginosa* grown on *C. albicans* WT compared to *adh1*Δ/Δ suggesting that a major portion of *P. aeruginosa* gene expression in co-culture was influenced by *C. albicans* ethanol production (**Fig. 1E** and **Supp. Dataset 1**). A similar trend was evident in *C. albicans* gene expression as approximately half of the DEGs between mono- and co-culture with *P. aeruginosa* were also DEGs between *C. albicans* WT and *C. albicans adh1*Δ/Δ from co-cultures (**Fig. 1F** and **Supp. Dataset 2**). Here, expression patterns of DEGs in both *P. aeruginosa* and *C. albicans* illustrated that ethanol played a defining role in *P. aeruginosa* – *C. albicans* interactions from the perspectives of both organisms.

### How fungal ethanol shapes co-culture transcriptomes: the *C. albicans* perspective

*C. albicans* Adh1 is responsible for reducing acetaldehyde to ethanol during fermentation, and we thus expected metabolic shifts between *C. albicans* WT and *adh1*Δ/Δ [29]. We identified DEGs between co-cultures of *C. albicans* WT and *adh1*Δ/Δ with *P. aeruginosa* and conducted KEGG [67–69] pathway over-representation analysis. The *C. albicans* KEGG pathway for fatty acid beta oxidation was over-represented in the DEGs and the genes it contained (e.g. *FAA2-1*, *FAA2-3*) were more highly expressed in *C. albicans* WT than in *C. albicans adh1*Δ/Δ (**Fig. 2, Supp. Dataset 3**). Since these pathways were not over-represented in DEGs between *C. albicans* WT and *C. albicans adh1*Δ/Δ in mono-culture, where there is no *P. aeruginosa*-produced 5-MPCA, these results are consistent with a previous report of a phenazine and other mitochondrial inhibitors increasing beta-oxidation in *C. albicans* WT as determined in metabolomics studies [70].

**Figure 2.**
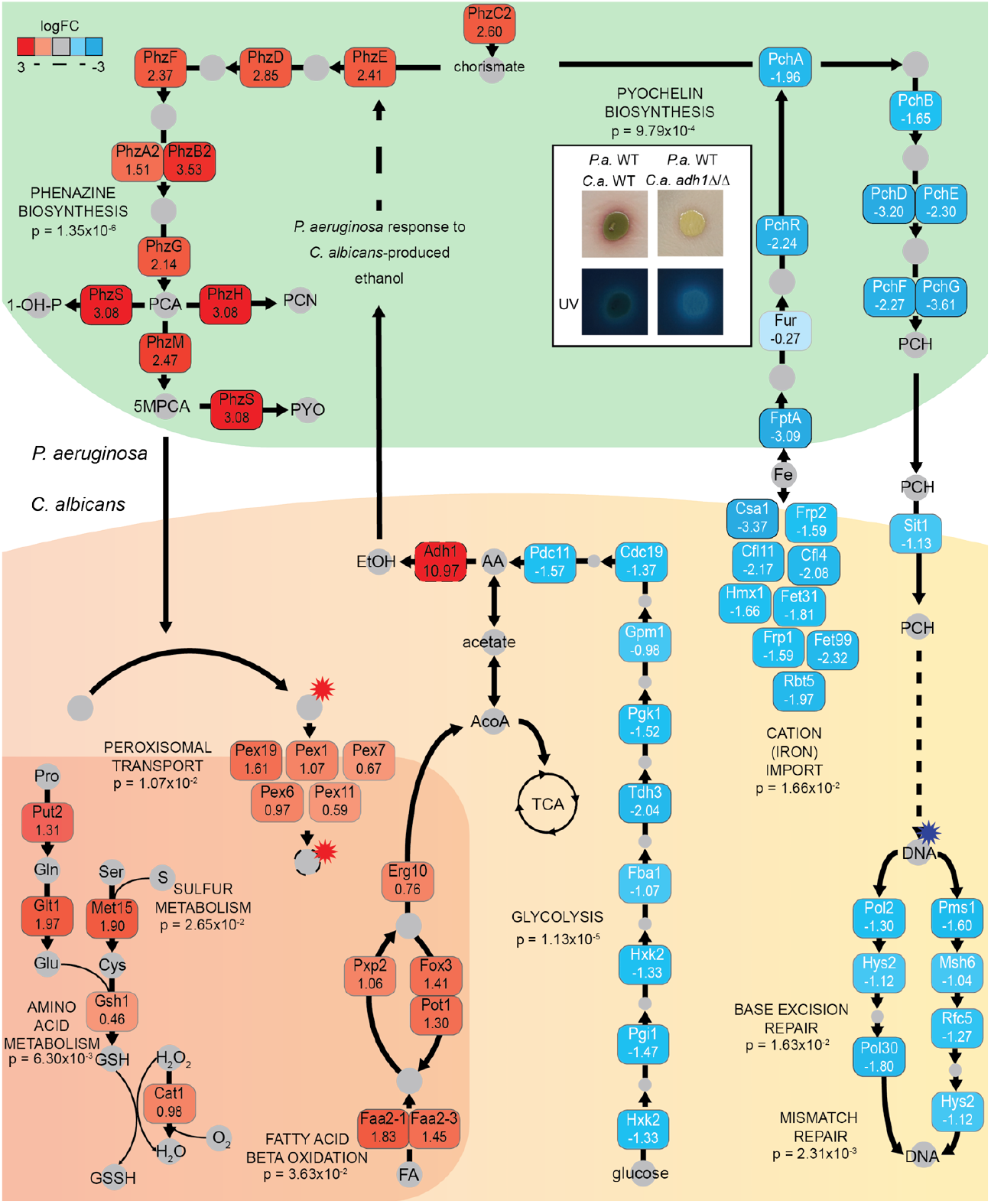
Pathways containing differentially expressed genes in *P. aeruginosa* and *C. albicans* between co-cultures of *P. aeruginosa* wild type (WT) with *C. albicans* WT or *adh1*Δ/Δ. DEGs of between *C. albicans* WT and *adh1*Δ/Δ from *P. aeruginosa* co-cultures contained over-representations of KEGG pathways for amino acid metabolism, sulfur (S) metabolism, peroxisomal transport, fatty acid beta-oxidation, glycolysis, cation (Fe) import, base excision and mismatch DNA repair. Red indicates higher expression in co-cultures with *C. albicans* WT and blue indicates higher expression in co-cultures with *C. albicans adh1*Δ/Δ. Values indicate log_2_fold-change. *C. albicans adh1*Δ/Δ has higher expression of glycolysis genes and the production of acetate, which is either secreted or enters into the citric acid cycle (TCA). *P. aeruginosa* DEGs from the same co-cultures were contained over-representations of KEGG pathways for phenazine (PCA, PCN, PYO, 1-OH-P, 5-MCPA) biosynthesis and pyochelin (PCH) biosynthesis pathways. Inset shows increase in siderophore-derived fluorescence of co-cultures of *P. aeruginosa* with *C. albicans adh1*Δ/Δ which is consistent with increased PCH production. P-values are from hypergeometric over-representation tests, FDR corrected.

Other KEGG pathways over-represented in DEGs in *C. albicans* from co-cultures of *C. albicans* WT with *P. aeruginosa* compared to *C. albicans adh1*Δ/Δ with *P. aeruginosa* were amino acid metabolism (e.g. *PUT2*, *GLT1*), sulfur metabolism (e.g. *MET15*) and peroxisomal transport (e.g. *PEX1*, *PEX19*) (**Fig. 2, Supp. Dataset 3**). These pathways converge on reactive oxygen species (ROS) mitigation (e.g. *GSH1*, *CAT1*) and, since previous reports have shown phenazines causing redox stress to neighboring fungi [25, 26], the upregulation of these pathways could have been due to ethanol-induced *P. aeruginosa* 5-MPCA production.

The KEGG pathway for glycolysis (e.g. *HXK2*, *PGI1*) was also over-represented in DEGs between *C. albicans* WT in co-culture with *P. aeruginosa and C. albicans adh1*Δ/Δ in co-culture with *P. aeruginosa,* but the genes within were more highly expressed in *C. albicans adh1*Δ/Δ compared to *C. albicans* WT, perhaps as metabolic compensation for the inability to ferment to ethanol (**Fig. 2, Supp. Dataset 3).** Similarly, there was also over-representation of the KEGG pathway for iron scavenging (e.g. *FRP1*, *FET99*) and these genes were again more highly expressed in *C. albicans adh1*Δ/Δ (**Fig. 2, Supp. Dataset 3).** Since the KEGG pathway for iron scavenging was not over-represented in DEGs between *C. albicans* WT and *C. albicans adh1*Δ/Δ in mono-culture (**Supp. Dataset 3)**, the increase in iron scavenging may have been due to a change in *P. aeruginosa* behavior that affected iron availability.

In co-cultures of *P. aeruginosa* with ethanol-deficient *C. albicans adh1*Δ/Δ, which did not promote 5-MPCA production*, C. albicans* also had higher expression of genes involved in DNA damage repair (e.g. *RBT5, CSA1*) (**Fig. 2**). While not a result of *P. aeruginosa* 5-MPCA, such damage may have been caused by another *P. aeruginosa* antagonistic factor. However, the KEGG pathways for DNA replication and repair were also over-represented in the DEGs between *C. albicans* and *adh1*Δ/Δ in mono-culture (**Supp. Dataset 3),** so DNA damage may be a native consequence of the loss of Adh1 in *C. albicans,* which is consistent with previous reports of an *adh1*Δ/Δ mutant having higher intracellular concentrations of the DNA-damaging metabolic intermediate methylglyoxal [71, 72].

### How fungal ethanol shapes co-culture transcriptomes: the *P. aeruginosa* **perspective**

We identified *P. aeruginosa* DEGs when grown on *C. albicans* WT compared to on *adh1*Δ/Δ. On *C. albicans* WT, *P. aeruginosa* upregulated genes involved in 5-MPCA biosynthesis including *phzM* and genes within both the *phz1* and *phz2* operons, which is consistent with differences in 5-MPCA formation between the two co-cultures (**Fig. 2**). While the last four genes of the *phz* operons have fewer than three SNPs between each gene pair and are thus not differentiated by alignment *phzA1, phzB1 and phzC1* have substantial enough differences in sequence from *phzA2, phzB2 and phzC2* respectively that the two operons can be distinguished, and we found that transcripts from both operons were more highly abundant by at least 2-fold when *P. aeruginosa* was grown in co-culture with *C. albicans* WT relative to with *adh1*Δ/Δ. While *phzS* and *phzH* are not required for 5-MPCA biosynthesis [24, 29] they have been reported to have coordinated expression with other phenazine genes [55, 73] and indeed we saw increases in their expression on *C. albicans* WT compared to *C. albicans adh1*Δ/Δ as well (**Fig. 2**).

We again conducted KEGG pathway over-representation analysis and found over-representation of three KEGG pathways in DEGs from *P. aeruginosa* grown in co-culture with *C. albicans* WT compared to with *C. albicans adh1*Δ/Δ: phenazine biosynthesis, quorum sensing (QS) and pyochelin biosynthesis (**Fig. 2, Supp. Dataset 3**). Over-representation of the phenazine biosynthesis pathway was expected based on the upregulation of the *phz* genes as described above. The identification of QS as an over-represented pathway was also not surprising in light of the known regulation of phenazine biosynthesis by QS in response to environmental cues [74, 75], including in *C. albicans* co-culture [27]. *P. aeruginosa* QS involves three major transcriptional regulators: LasR [76], PqsR [77] and RhlR [76]. RhlR and PqsR were necessary for phenazine production (**Fig. S1B**). Although Δ*lasR* appeared to produce less 5-MPCA than *P. aeruginosa* WT on *C. albicans* WT, consistent with previous data, it produced an abundance of the blue-green phenazine pyocyanin, which is a 5-MPCA-derivative [18]. Upon examining the expression of gene targets for these transcription factors (**Supp. Dataset 6**), we found heterogenous expression patterns inconsistent with canonical, cell-density regulated QS but reconcilable with activation of a subset of QS regulated genes that includes phenazine biosynthesis genes (**Fig. S1C,D)**.

The third over-represented KEGG pathway was that for the biosynthesis of pyochelin, a siderophore [51]. Expressly, genes involved in pyochelin biosynthesis, import and regulation were lower in *P. aeruginosa* on *C. albicans* WT compared to on *C. albicans adh1*Δ/Δ. Pyochelin and another siderophore, pyoverdine, are fluorescent, and we supported the RNA-Seq data by showing increased *P. aeruginosa*-derived fluorescence on *C. albicans adh1*Δ/Δ compared to on *C. albicans* WT (**Fig. 2**, inset). The over-representation of low iron responsive genes in both *P. aeruginosa* and *C. albicans* demonstrated that the organisms were experiencing simultaneous iron limitation. Taken together these data are consistent with 1) *C. albicans* ethanol production stimulated *P. aeruginosa* antagonistic 5-MPCA production which affected *C. albicans* metabolism and ROS stress pathways and 2) increased glycolysis in *C. albicans adh1*Δ/Δ that compensated for the inability to ferment coincided with a competition for iron.

### eADAGE analysis of *P. aeruginosa* transcriptome revealed additional pathways differentially active in response to ethanol in co-culture with *C. albicans*

While the analysis of DEGs and the KEGG pathways over-represented therein provided insight into two key *P. aeruginosa* – *C. albicans* interactions, only 19 of the 120 *P. aeruginosa* DEGs (|ogFC > 2, FDR < 0.05) fell within the three statistically over-represented KEGG pathways: QS (orange bar), phenazine biosynthesis (red bar) and pyochelin biosynthesis (blue bar) (**Fig. 3A**). To look for additional processes that were affected by *C. albicans* ethanol production, we further identified patterns in the RNA-Seq data using eADAGE, a denoising autoencoder-based tool [53, 58, 64]. In eADAGE analysis, the activities of previously-defined gene expression signatures are calculated as a weighted sum of normalized gene expression values (TPM) where gene weights are unique to each signature [64]. The eADAGE signatures were derived irrespective of human curation, which allowed for the examination of gene sets with coherent expression patterns but no annotation to date. The eADAGE-transformed *P. aeruginosa* – *C. albicans* co-culture signature activity profiles had a higher clustering coefficient (CC) by condition than was observed by normalized gene expression profiles; the CC for differentially active eADAGE signatures (DASs) was 0.68 compared to 0.38 for gene expression data; randomized data had CC values less than 0.14 (**Table S1**). The higher CC after eADAGE signature transformation indicated that biological information was retained and signals that differentiated sample types may be more clear at the pathway level. Using eADAGE, we found 48 DASs in *P. aeruginosa* grown on *C. albicans* WT versus *P. aeruginosa* grown on *C. albicans adh1*Δ/Δ (**Supp. Dataset 3**).

**Figure 3.**
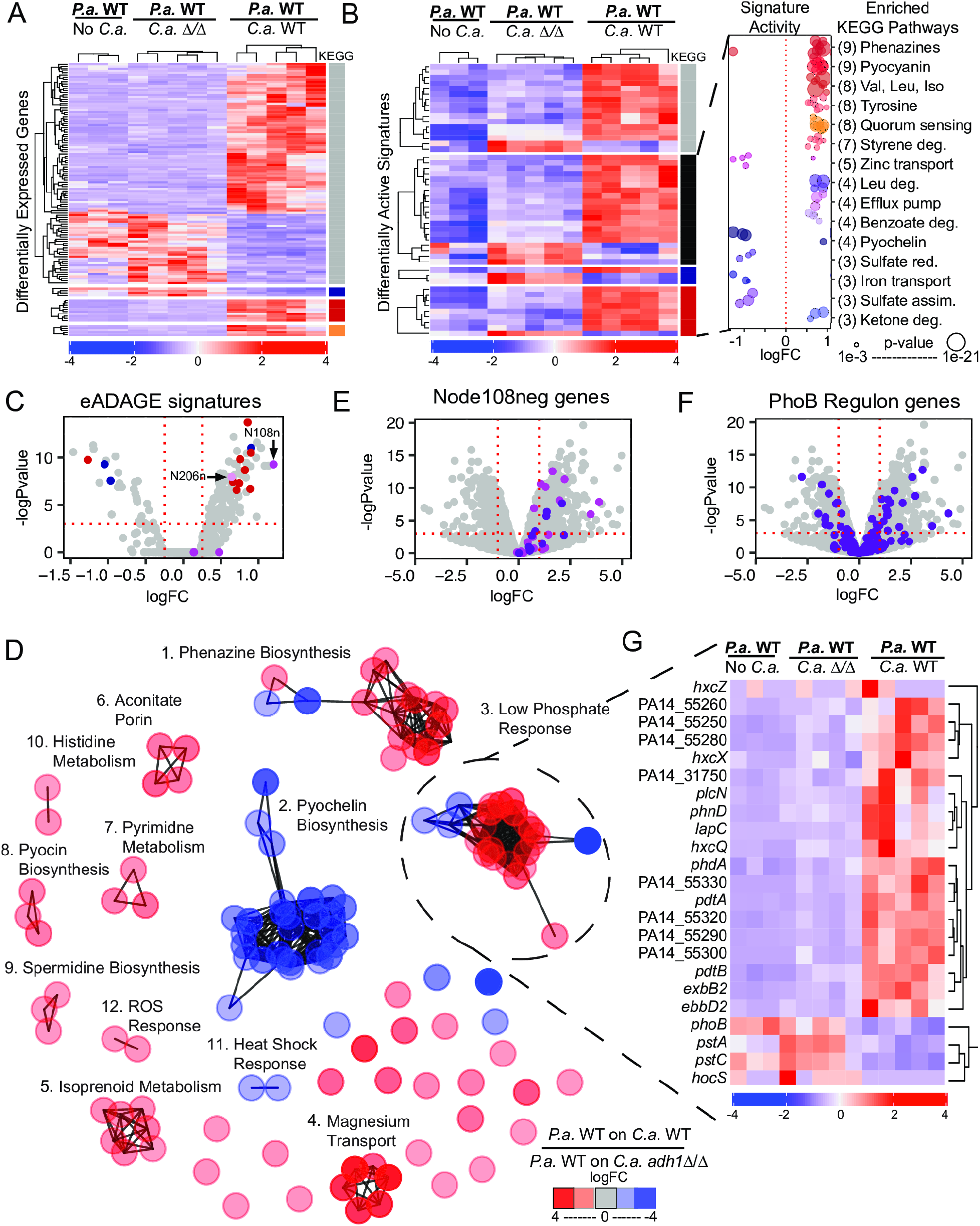
eADAGE analysis reveals a subset of the Pho regulon upregulated in *P. aeruginosa* (*P.a.*) grown on *C. albicans* (*C.a.*) WT compared to that on *C.a*. *adh1*Δ/Δ. A) Differentially expressed genes (DEGs) between *P.a.* grown alone or on *C.a.* WT and *C.a. adh1*Δ/Δ for 24 h. Genes that fell within over-represented KEGG pathways (quorum sensing (orange bar), phenazine biosynthesis (red bar) and pyochelin biosynthesis (blue bar)) are indicated. Most DEGs do not belong to any of the three pathways (grey bar). B) Differentially active eADAGE signatures (DASs) for the same samples shown in A. Signatures in which genes annotated as being involved in phenazine biosynthesis (red bar), pyochelin biosynthesis (blue bar), or other KEGG pathways (black bar) are overrepresented are indicated. Signatures that are not over-represented an any KEGG pathways are indicated by the grey bar. Inset shows the fold-change for the expression all of the KEGG pathways that are over-represented among the DASs (# of DASs per KEGG pathway in parentheses); over-representation p-value shown as circle (Supp. Dataset 3). C) DASs with increased activity in transcriptome comparisons of *P.a*. grown on *C.a*. WT compared to on *C.a*. *adh1*Δ/Δ. In addition to DASs with over-representations of pyochelin (blue dots) and phenazine (red dots) biosynthesis, others over-represent the Pho regulon (Node108n, purple) or contain ethanol catabolism genes (N206n, pink). D) The eADAGE signature with the highest increase in activity, Node108neg (N108n, purple), contains many genes with increased expression though not all met the criterion of DEGs individually (logFC > 1,,FDR < 0.05). E) DEGs in *P.a*. grown on *C.a*. WT compared to on *C.a*. *adh1*Δ/Δ with expression levels of PhoB-regulated genes (dark purple) highlighted. F) Network analysis of DEGs suggest groups of DEGs have correlated patterns across eADAGE: phenazine biosynthesis (1) is inversely expressed with the low iron response (2) and coordinately upregulated with the low phosphate response (3) upon exposure to ethanol in co-culture. Other cliques of DEGs participate in shared biological pathways. See table 2 for descriptions of all cliques. G) The Pho Clique (3) contains two clades of DEGs with opposing expression patterns between *P.a.* grown on *C.a.* WT and *C.a*. *adh1*Δ/Δ.

**Table 1.** Phenotypes for 5MPCA-accumulation in co-culture *C. albicans* WT by *P. aeruginosa* strains and ethanol-induced alkaline phosphatase (AP) activity in monoculture. AP activity was visualized in colonies on agar containing by BCIP. *P. aeruginosa* mutants were defective in ppGpp-dependent AlgU and DksA signaling, ethanol catabolism, the kinase KinB and sigma factor VreI, which are known to influence PhoB activity, acetyl-phosphate metabolism and ethanol catabolism.

| Genotype | *P.a.*<br>5MPCA on<br>*C. albicans*<br>WT | *P.a.* AP<br>induction<br>by<br>ethanol |
| --- | --- | --- |
| WT | Yes | Yes |
| $\Delta phoB$ | No | No |
| $\Delta algU$ | Yes | Yes |
| $\Delta dksA$ | Yes | Yes |
| $\Delta relA$ | Yes | Yes |
| $\Delta relA\Delta spoT$ | Yes | Yes |
| $\Delta kinB$ | No | Yes |
| $\Delta kinB+kinB$ | Yes | Yes |
| $\Delta mucB$ | Yes | Yes |
| $\Delta vrel$ | | Yes |
| $\Delta vreR$ | | Yes |
| $\Delta vreA$ | | Yes |
| $\Delta vrel\Delta phoB$ | | No |
| $\Delta exaA$ | No | No |
| $\Delta exaA+exaA$ | Yes | Yes |
| $acsA::TnM$ | Yes | No |
| $\Delta ackA$ | Yes | Yes |
| $\Delta ackA\Delta pta$ | Yes | Yes |
| $\Delta ackA\Delta pta\Delta phz$ | No | Yes |

As predicted by the DEG analysis (**Fig. 3A**), there were multiple DASs in which phenazine biosynthesis and pyochelin biosynthesis KEGG pathways were over-represented (**Fig. 3B**, red and blue bars respectively). The detection of multiple signatures enriched in these pathways was expected based on the presence of similar signatures that are redundant in some contexts but discriminating in others [53]. There were 32 DASs (**Fig. 3B**, black bar) with over-representations of 7 additional pathways (all pathways over-represented in DASs shown in inset) and 16 DASs in which no pathway was over-represented (**Fig. 3B**, grey bar), perhaps representing novel transcriptional signals. As expected based on the larger differences in gene expression for phenazine biosynthetic genes, multiple signatures that contained over-representations of phenazine biosynthesis (annotated as phenazine or pyocyanin) (red) had increased activity in *P. aeruginosa* grown on *C. albicans* WT compared to *P. aeruginosa* grown on *C. albicans adh1*Δ/Δ, and together contained over-representations of pyochelin biosynthesis (annotated as siderophore or iron transport) (blue) had decreased activity in *P. aeruginosa* grown on *C. albicans* WT compared to *P. aeruginosa* grown on *C. albicans adh1*Δ/Δ. Other pathways over-represented in DASs included amino acid metabolism, styrene metabolism and zinc uptake (**Fig. 3B** inset, **Supp. Dataset S4**). Notably, one differentially active signature, Node206neg (N206n), contained ethanol catabolism genes (**Fig. 3C**, pink dot). The signature with the largest increase in activity, Node108neg (N108n) (**Fig. 3C**, violet dot) was not enriched in any KEGG pathways. However, we had previously identified Node108neg as significantly more active in low phosphate media than phosphate replete media across the compendium of gene expression on which eADAGE was trained [53]. Therefore, upon identifying Node108neg as the most activated eADAGE signature in response to *C. albicans* ethanol production in co-culture, we further investigated the genes within Node108neg and their connection to the *P. aeruginosa* low phosphate response.

### eADAGE analysis suggests *P. aeruginosa* PhoB up-regulated genes in response to ***C. albicans* ethanol production**

In the most upregulated eADAGE signature, Node108neg, the set of PhoB-regulated genes PhoB (i.e. the PhoB regulon) was significantly over-represented (hypergeometric test: p = 5.9×10^-9^). The PhoB regulon has been extensively defined through rigorous experimental methods including mutant transcriptomics, motif analysis and chromatin immunoprecipitation assays [55]. Of the 32 genes in Node108neg, 11 were also in the PhoB regulon (**Fig 3D**) and they all increased in expression in *P. aeruginosa* grown on *C. albicans* WT compared to *adh1*Δ/Δ, but the Pho regulon defined in Bielecki *et al*. was heterogeneously expressed overall (**Fig. 3E**). We examined DEGs in the context of signatures more closely in order understand how changes in eADAGE signature activities embodied the *P. aeruginosa* response to *C. albicans*-produced ethanol and whether *P. aeruginosa* gene expression changes between growth on *C. albicans* WT and *adh1*Δ/Δ signaled a low phosphate response.

We visualized relationships among DEGs in the eADAGE gene-gene network. The full gene-gene network consists of the 5,549 *P. aeruginosa* genes used to create the eADAGE model as vertices with similarities in transcriptional patterns as weighted edges (shorter edges represent higher Pearson correlations between gene weights across all signatures in the eADAGE model) [53, 58, 64]. Here we show strongly DEGs (logFC > 2, p-value < 0.01) between *P. aeruginosa* grown on *C. albicans* WT and *adh1*Δ/Δ connected by edges whose weight is drawn from the full gene-gene network (edge cutoff ± 0.5) (**Fig. 3F**). DEGs fell into cliques (**Table S2**) when visualized as a sub-network within the eADAGE gene-gene network (**Fig. 3D**). The three largest cliques contained genes relevant to the biological processes of phenazine biosynthesis (clique 1, 17 genes), pyochelin biosynthesis (clique 2, 29 genes), and the low phosphate response (clique 3, 23 genes) (**Fig. 3F**). Other cliques contained genes involved in isoprenoid catabolism (clique 4), magnesium flux across the membrane (clique 5), aconitate porins (clique 6), pyrimidine metabolism (clique 7), pyocin biosynthesis (clique 8), spermidine biosynthesis (clique 9), histidine metabolism (clique 10), the heat shock response (clique 11), and the ROS stress response (clique 12). Most notably, many genes within clique 3, which were related to the low phosphate response, were also in Node108neg.

Genes within clique 3 clustered into two groups by gene expression: four were more highly expressed in the *P. aeruginosa* grown on *C. albicans adh1*Δ/Δ and 19 genes were more highly expressed in *P. aeruginosa* grown on *C. albicans* WT, including those in Node108neg (**Fig. 3G**). Notably, both groups of genes are regulated by PhoB, but are in different operons. The group of *P. aeruginosa* genes that were more highly expressed when grown on *C. albicans adh1*Δ/Δ fall within the neighboring operons *phoBR* and *pstABC*. The group of *P. aeruginosa* genes that were more highly expressed when grown on *C. albicans* WT belong to the consecutive operons that encode the Hxz type II secretion system (PA14_55450, PA14_55460) and its substrate enzyme the phosphatase *lapC* [78]. The latter group of genes also contained genes involved in TonB-dependent transport (*exbB2*, *exbD2*) as well as the phosphatases *phoA*, and the phospholipases *plcN*. All of these genes are canonically co-regulated by PhoB and usually have positively correlated expression patterns but it appeared as if PhoB was selectively promoting the expression of only a subset of its regulon. We demonstrated that the non-canonical PhoB response identified by eADAGE was biologically meaningful through genetic, biochemical and phenotypic experiments described below.

### The *P. aeruginosa* low phosphate response was activated in response to *C. albicans* ethanol production

eADAGE analysis of the co-culture RNA-Seq data found higher levels of some PhoB-regulated transcripts in *P. aeruginosa* co-cultured with *C. albicans* WT than in *P. aeruginosa* co-cultured with *C. albicans adh1*Δ/Δ. We confirmed this result using NanoString, a multiplex RNA analysis method, to measure the mRNA levels of representative PhoB-regulated genes. The data shown are normalized to six house keeping genes as described in the Methods section. Like in the RNA-Seq data, there was a split in the expression of PhoB-regulated genes: phosphate transport-associated genes *phoR*, *phoB*, *pstA* and *phoU* did not increase in expression (**Fig. 4A**, bottom section) but those encoding phosphate scavenging enzymes did: glycosyl transferase PA14_53380, glycerophosphoryl diester phosphodiesterase *glpQ*, TonB-dependent transport protein *exbD2,* phosphonate transporter *phnD* and alkaline phosphatase *phoA*. As well as genes for phenazine biosynthesis (**Fig. 4A**, top two sections). Additionally, ethanol catabolism genes *exaA* and *exaB* showed PhoB-independent increases in expression (**Fig. 4A**, third section).

**Figure 4.**
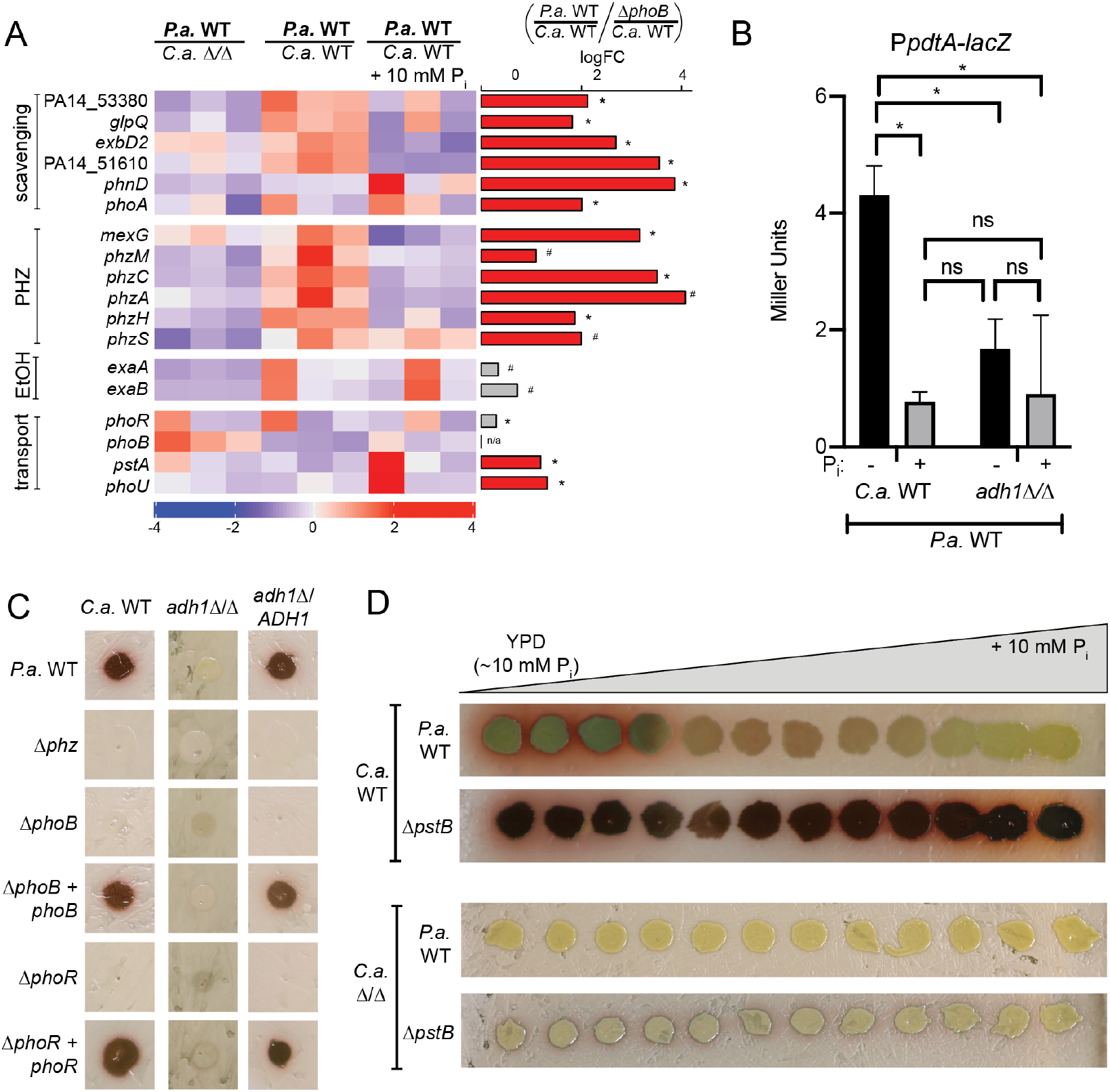
*C. albicans* (*C.a.*) WT induced PhoB-regulated genes in *P. aeruginosa* (*P*.*a*.) compared to *C.a. adh1*Δ/Δ leading to 5-MPCA production as indicated by red pigment formation. A) Expression of *P.a.* genes involved in phosphate scavenging, phenazine biosynthesis (PHZ), ethanol catabolism (EtOH) and inorganic phosphate transport were measured by NanoString (codesetPAV5) from cells grown with *C.a. adh1*Δ/Δ, *C.a.* WT or *C.a.* WT grown on medium with additional 10 mM phosphate. Expression values are normalized to loading controls and housekeeping genes as described in methods. Values are z-scored, scaled by gene. Right-hand barplot shows logFC between *P.a.* WT and *P.a*. Δ*phoB* on *C.a.* WT. The bar is colored red if expression is PhoB-dependent (logFC *P.a*. WT / *P.a.* Δ*phoB* > 1, FDR < 0.05, else grey). * = FDR < 0.05, # = FDR > 0.05. B) Beta-galactosidase activity indicative of expression of a *pdtA*-*lacZ* promoter fusion in *P.a.* WT in *P.a* grown with *C.a.* WT or *C.a*. *adh1*Δ/Δ in the absence or presence of P_i_ supplementation, *,p<0.05 by ANOVA (n = 3). C) Red 5-MPCA derivatives produced in co-culture by *P.a* WT, Δ*phoB*, Δ*phoR*, and their complemented derivatives on *C.a.* WT, *adh1*Δ/Δ, and *adh1*Δ/*ADH1*. D) Red 5-MPCA-derivatives produced by *P.a.* WT and *P.a.* Δ*pstB* over a gradient of phosphate concentrations. *P.a.* Δ*pstB* has constitutive PhoB activity. Conversely, *P.a.* WT did not produce 5-MPCA on *Ca. adh1*Δ/Δ at any phosphate concentration, but *P.a.* Δ*pstB* induced a small amount of red pigment independent of the phosphate concentration.

We assessed PhoB activity in co-culture using phosphate supplementation, the native suppressor of PhoB. The addition of 10 mM potassium phosphate to the medium underlying the co-cultures resulted in a decrease in the expression levels of PhoB regulated genes (**Fig. 4A**, top section), including phenazine genes (**Fig. 4A**, second section). Visualization of transcript levels by sample revealed some heterogeneity which we predicted was indicative of a dynamic response as both species grow. To additionally confirm the expression of these genes was PhoB-dependent we included Δ*phoB* in co-culture with *C. albicans* WT. The histogram to the right shows the mean signal of PhoB-dependence (*P. aeruginosa* WT on *C. albicans* WT / *P. aeruginosa* Δ*phoB* on *C. albicans* WT) and indicates that expression of expected PhoB targets but not ethanol catabolism genes was PhoB-dependent.

Complementing transcript abundance data, promoter activity in *P. aeruginosa* WT decreased in response to 10 mM phosphate in co-culture with *C. albicans* WT as measured by promoter fusion assay of the PhoB target *pdtA* [41], **Fig. 4B**). These data suggested that in co-culture, the bioavailability of phosphate modulated PhoB activity which may have affected co-culture interactions.

### PhoB was necessary for 5-MPCA-derived red pigment accumulation in *P. aeruginosa* – *C. albicans* co-culture

Given the dependence on PhoB for the expression of phenazine biosynthesis genes, we determined if PhoB was necessary for the accumulation of 5-MPCA-derived red pigment in co-culture with *C. albicans* WT. *P. aeruginosa* Δ*phoB* did not support accumulation of the 5-MPCA-derivative as indicated by the lack of red pigment in co-culture with ethanol-producing *C. albicans* WT, and this was restored by chromosomal complementation with a wild-type copy of *phoB* in *P. aeruginosa,* provided *C. albicans* had a functional *ADH1* gene (**Fig. 4C**). Phenazine production was also dependent on PhoR, the known regulatory kinase of PhoB, and the Δ*phoR* phenotype was complemented by a wild-type copy of *phoR* expressed on an extrachromosomal plasmid. We further demonstrated that PhoB activity is required for 5-MPCA production as phosphate supplementation led to a decrease in 5-MPCA-derived red pigment in *P. aeruginosa* WT co-culture with *C. albicans* WT as seen across a gradient plate (**Fig. 4D**, top). PhoB activity was sufficient to overcome suppression by phosphate supplementation as a mutant lacking the phosphate transport ATPase *pstB* with constitutively active PhoB, continued to produce 5-MPCA-derived red pigment despite the addition of phosphate (**Fig. 4E**, second down). As expected, *P. aeruginosa* WT did not form any 5-MPCA-derived pigment in co-culture with *C. albicans adh1*Δ/Δ at any phosphate concentration tested (**Fig. 4E**, third down). *P. aeruginosa* Δ*pstB* grown on *adh1*Δ/Δ only slightly rescued 5-MPCA-derived red pigment formation (**Fig 4**. **D**, bottom). This suggested that, while phosphate levels contributed to the control of PhoB activity, *C. albicans* Adh1 activity provided an additional stimulus that created conditions conducive to PhoB-regulated *P. aeruginosa* antifungal phenazine production.

### Ethanol was sufficient to activate PhoB at intermediate phosphate concentrations

To determine if ethanol was sufficient to stimulate PhoB in *P. aeruginosa*, we monitored PhoB-regulated alkaline phosphatase (AP) activity. AP activity can be monitored by the conversion of 5-bromo-4-chloro-3-indolylphosphate (BCIP) into a colorimetric substrate according to published methods [79–81]. On MOPS (3-morpholinopropane-1-sulfonic acid) buffered minimal medium after growth for 16 h at 37°C, in *P. aeruginosa* Δ*phz* where all blue coloration can be attributable to BCIP conversion and not phenazines, AP was detected up to approximately 0.55 mM phosphate in the absence of ethanol (**Fig. 5A**). With the addition of 1% ethanol to the medium, *P. aeruginosa* showed AP activity at higher phosphate concentrations, up to approximately 0.73 mM phosphate (**Fig. 5A**). This induction was also seen in *P. aeruginosa* WT but not Δ*phoB* at 0.7 mM phosphate on single concentrations plates, and was restored upon complementation with a WT copy of *phoB* (**Fig. 5B**). Quantification of AP activity under these conditions showed a 4-fold increase in the presence of 1% ethanol for WT *P. aeruginosa*, only trace AP activity for *P. aeruginosa* Δ*phoB* and hyper activity in Δ*pstB* (**Fig. 5C**).

**Figure 5.**
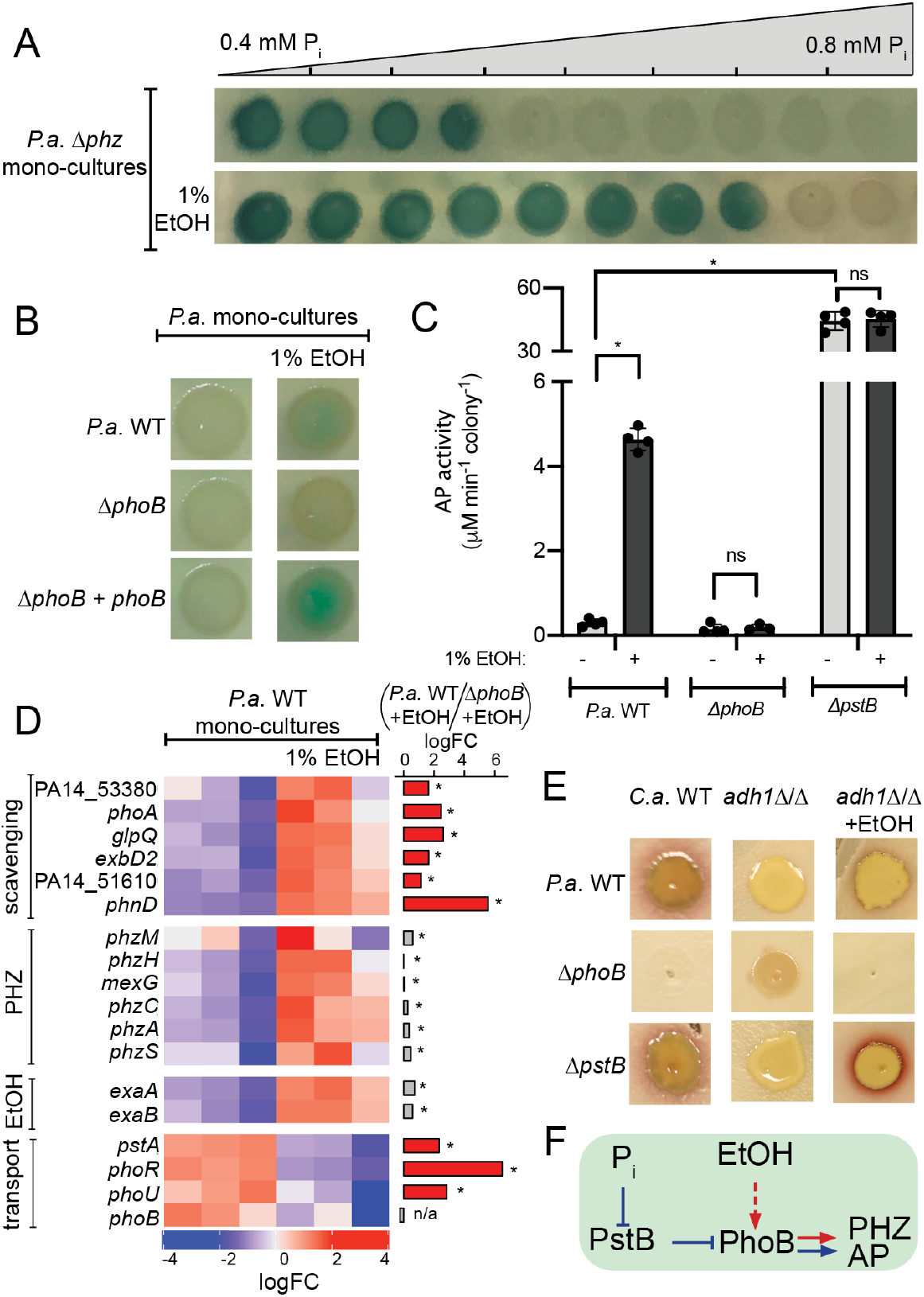
Ethanol (EtOH) induced PhoB activity in *P. aeruginosa* (*P.a.*) mono-culture. A) Alkaline phosphatase (AP) activity visualized by blue color derived from cleavage of BCIP in MOPS medium with a gradient of phosphate in the absence and presence of ethanol. The *P.a.* Δ*phz* strain was used to eliminate color differences due to phenazine production. B) AP activity in the absence and presence of ethanol was visualized with BCIP added to MOPS agar (0.7 mM phosphate) for *P.a.* wild type, Δ*phoB* and the Δ*phoB* mutant complemented with a wild-type copy of *phoB* integrated at the native locus. C) AP activity in cells from colony biofilms grown as in B was measured using the colorimetric substrate pNPP for *P.a.* WT, Δ*phoB*, and Δ*pstB*, a strain with constitutive PhoB activity. *,p<0.01 by ANOVA (n ≥ 3). D) Transcripts within the PhoB regulon involved in phosphate scavenging and inorganic phosphate transport and genes involved in phenazine production (PHZ) and ethanol catabolism (EtOH) were measured in cells grown in the absence and presence of ethanol by Nanostring (codeset PAV5). PhoB-regulated genes increased in expression (top section) and ethanol catabolism genes (third section) increase in expression and others decrease (bottom section). Expression values are normalized to loading controls and housekeeping genes as described in methods. Values are scaled by gene. Right-hand barplot shows logFC between *P.a.* WT and *P.a*. Δ*phoB* on MOPS+1%EtOH. The bar is colored red if expression is PhoB-dependent (logFC *P.a*. WT / *P.a.* Δ*phoB* > 1, FDR < 0.05, else grey). *,FDR < 0.05, #,FDR > 0.05. E) PhoB activity and ethanol are both for 5-MPCA production in response to *C.a.* ethanol. P.a. WT, Δ*phoB* and Δ*pstB* were grown on C.a. WT, or C.a. adh1Δ/Δ in the absence or presence of exogenous ethanol. 5-MPCA production was rescued by the addition of 1% ethanol to a co-culture of *P.a.* Δ*pstB* and, to a lesser extent, *P.a.* WT on *C.a. adh1*Δ/Δ. F) Phosphate (P_i_) and ethanol (EtOH) are additive stimuli that promote PhoB-dependent expression of AP and phenazine biosynthesis (PHZ).

To determine if PhoB was acting upon the same targets under ethanol supplementation in mono-culture as shown for co-culture, we monitored the expression of PhoB targets using the same Nanostring codeset described earlier. *P. aeruginosa* WT and Δ*phoB* were grown in mono-culture on MOPS minimal medium with 0.7 mM phosphate with and without 1% ethanol. The same subset of PhoB-regulated genes indicated in co-culture expression analysis increased in *P. aeruginosa* WT upon ethanol supplementation: *phoA*, an alkaline phosphatase; *phnD,* a phosphonate transporter; *glpQ*, a glycerophosphoryl diester phosphodiesterase (**Fig. 5D**, top section). In mono-culture, when ethanol was supplemented into the medium, we saw an increase in phenazine biosynthesis genes however, unlike in co-culture, the magnitudes of their logFC between *P. aeruginosa* WT and Δ*phoB* was not as high as those of other PhoB-regulated genes suggesting their expression was also stimulated by PhoB-independent factors (**Fig. 5D**, second panel). Interestingly, the set of PhoB-regulated genes whose expression was heterogenous and did not trend upward in co-culture had expression that trended downward in mono-culture upon ethanol supplementation (**Fig. 5D**, bottom panel). Stimulation of PhoB by ethanol in mono-culture meant that ethanol could have been one of the stimuli that activated PhoB in co-culture.

### Low phosphate and ethanol were additive stimulants for PhoB-mediated red pigment formation

We tested whether ethanol stimulation and canonical PhoB de-repression (Δ*pstB*) had additive effects on 5-MPCA-derived red pigment formation in *P. aeruginosa – C. albicans* co-culture. On non-ethanol producing *C. albicans adh1*Δ/Δ, the constitutive PhoB activity in *P. aeruginosa* Δ*pstB* was not sufficient to stimulate red pigment as that on ethanol-producing *C. albicans* WT (**Fig. 5E**). Strikingly, the addition of 1% ethanol to *C. albicans* a*dh1*Δ/Δ lawns increased red pigment formation in *P. aeruginosa* Δ*pstB* but not in *P. aeruginosa* WT or Δ*phoB* (**Fig. 5E**). This signified that PhoB activation both through the canonical signaling pathway and by means of ethanol stimulation were necessary and only combined were sufficient for 5-MPCA-derived red pigment formation in co-culture (proposed model in **Fig. 5F**).

In light of the recent characterization of the *P. aeruginosa* response to exogenous ethanol [31] that showed ppGpp, synthesized by RelA and SpoT, DksA, and AlgU mounted a transcriptional response to ethanol, we assessed their role in ethanol-induced PhoB activation in co-culture by monitoring red pigment accumulation and in mono-culture via BCIP assay. We determined that these genes were not necessary to induce PhoB activity in response to ethanol (**Table 1**). We also ruled out roles for known mechanisms of alternative PhoB activation including contributions of the non-canonical histidine kinase KinB in both co- and mono-culture [53] and the extra-cytoplasmic function (ECF) sigma factor VreI in monoculture, but further investigation is required to determine if VreI played a role in regulating 5-MPCA production in co-culture [37, 41] (**Table 1**).

We tested the role of ethanol catabolism in PhoB activation in both mono- and co-culture. Mutants in the ExaA-dependent pathway for ethanol catabolism through acetate including *exaA* and *acsA* did not show increases in AP activity in response to 1% ethanol (**Table 1**) indicating that ethanol catabolism was essential for PhoB activation in mono-culture. We hypothesized that ethanol catabolism led to increased levels of acetyl phosphate, a non-canonical phosphate donor of transcription factors including PhoB, but neither acetyl phosphate biosynthesis mediated by AckA nor catabolism by Pta was necessary for ethanol-induced PhoB activity (**Table 1**) [82–85]. Mutants defective in the ExaA-dependent ethanol catabolic pathway showed only weak 5-MPCA production on *C. albicans* lawns and the phenotype could be complemented (**Table 1**). Together these data suggest that ethanol catabolism plays a role in PhoB activation, and that additional pathways for ethanol catabolism may be present in co-culture conditions. Further investigation is required to determine the mechanism for ethanol induction of PhoB activity, but these results demonstrate ethanol stimulation and PhoB activation participated in non-linear but intersecting pathways.

### The *C. albicans* low phosphate response was more active in *adh1*Δ/Δ than in WT

Given the differences in *P. aeruginosa* PhoB activity in co-culture with *C. albicans* WT compared *adh1*Δ/Δ, we sought to understand if *C. albicans* also experienced differences in phosphate availability by examining its low phosphate response. In *C. albicans*, low phosphate induces activity of the transcription factor Pho4 which regulates genes involved in phosphate acquisition as well as the homeostasis of cations (e.g. iron), tolerance of stresses including ROS and arsenic, and fitness in murine models [86–90]. Pho4 regulates 133 genes that were identified using transcriptomics as differentially expressed both between *C. albicans* WT and *pho4*Δ/Δ in low phosphate and between *C. albicans* WT in high and low phosphate [86] (see **Supp. Dataset 4** for gene list). We found that Pho4-regulated genes were over-represented in DEGs between co-cultures of *C. albicans* WT and *C. albicans adh1*Δ/Δ (logFC > 1, p < 0.05) (hypergeometric test p = 1.9×10^-3^). Of the top ten most differentially expressed genes between *C. albicans* WT and *pho4*Δ/Δ in low phosphate determined by Ikeh *et al.* [86], seven were also strongly differentially expressed (logFC > 2) in co-cultures of *C. albicans* WT with *P. aeruginosa* compared to *C. albicans adh1*Δ/Δ with *P. aeruginosa* including a secreted phospholipase (*PHO100*) and two secreted phosphatases (*PHO112* and *PHO113*) (**Fig. 6A**, black bars). Given the strong differential expression of these phosphatases, we assessed phosphatase activity using BCIP supplementation as done for *P. aeruginosa*. Consistent with the transcriptional data, higher phosphatase activity was observed in *C. albicans adh1*Δ/Δ compared to *C. albicans* WT in the presence (**Fig. 6B**) of *P. aeruginosa*.

**Figure 6.**
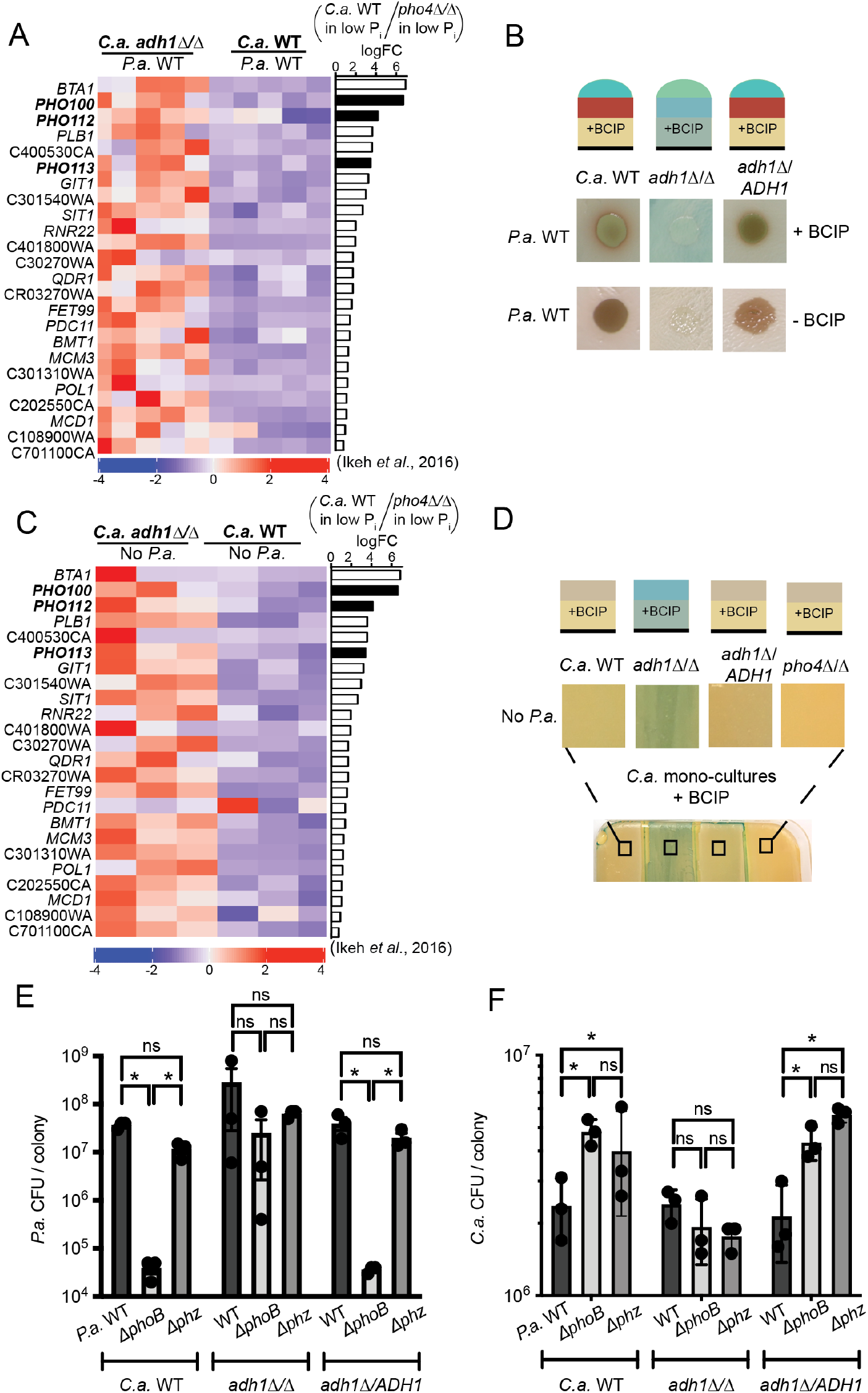
The *C. albicans* (*C.a*.) *adh1*Δ/Δ has increased expression of the Pho4-mediated low phosphate response in co-culture that is inversely correlates with PhoB activity in *P. aeruginosa* (*P.a*.). A) Previously characterized Pho4-regulated genes [86] including a phospholipase and phosphatases (black bars) were more highly expressed in *C.a*. *adh1*Δ/Δ than *C.a.* WT in *P.a*. co-cultures (data shown as z-scores of TPM). Pho4-dependence is shown in the right-hand barplot as log_2_FC *C.a*. WT/*C.a. pho4*Δ/Δ using data from [86]. B) Analysis of phosphatase activity in *C.a.* WT, *C.a. adh1*Δ/Δ, the complemented strain *adh1*Δ/*ADH1* or *pho4*Δ/Δ using the colorimetric phosphatase BCIP substrate in agar. More phosphatase activity was observed in *C.a. adh1*Δ/Δ than in strains with *ADH1* in in co-culture with *P.a*. C) The same Pho4-regulated genes as shown in A were also more highly expressed in *C.a. adh1*Δ/Δ than *C.a.* WT in mono-cultures. Right-hand barplot shows Pho4-dependence as in A. D) More phosphatase activity was observed in *C.a. adh1*Δ/Δ than in strains with *ADH1* in in mono-culture.. As predicted, phosphatase activity is not evident in the *C.a*. *pho4*Δ/Δ strain. Phosphatase activity visualized via BCIP as in B. E) Number of CFUs of *P.a*. WT, Δ*phoB* or Δ*phz* after co-culture for 72 h with *C.a.* WT, *C.a. adh1*Δ/Δ *C.a. adh1*Δ/*ADH1*. *P.a.* Δ*phoB* and Δ*phz* had significantly fewer CFUs on *C.a.* strains with high ethanol production (WT and *adh1*Δ/*ADH1*), but not *C.a. adh1*Δ/Δ. F) In the same samples analyzed in A, *C.a*. CFU formation was assessed. *C.a.* WT or *C.a. adh1*Δ/*ADH1* strains had increased fitness in-culture with *P.a*. Δ*phoB* or *P.a*. Δ*phz* compared to with *P.a*. WT. For *C.a. adh1*Δ/Δ, there were no differences in CFU formation when co-cultured with *P.a*. WT, Δ*phoB* or Δ*phz*.

To determine if the activation of the *C. albicans* low phosphate response in *adh1*Δ/Δ in co-culture was a consequence of competition for phosphate with *P. aeruginosa*, we examined *C. albicans* gene expression and phosphatase activity in mono-culture. The same Pho4-regulated genes that were DEGs in co-culture were also DEGs between *C. albicans* WT and *C. albicans adh1*Δ/Δ even in the absence of *P. aeruginosa* (**Fig. 6C**), and higher phosphatase activity was evident in *C. albicans adh1*Δ/Δ compared to *C. albicans* WT or the complemented strain *adh1*Δ/*ADH1* in mono-culture by BCIP assay (**Fig 6D**). As a control, we assayed *C. albicans pho4*Δ/Δ. The low amounts of phosphatase activity seen for *C. albicans* WT and *adh1*Δ/*ADH1* was distinguishable from the absence of phosphatase in *pho4*Δ/Δ indicating that that phosphatase activity was Pho4-dependent (**Fig 6D**). The high Pho4 response in *C. albicans adh1*Δ/Δ, evident even in mono-culture, suggested that Adh1 activity impacts *C. albicans* phosphate access, requirements or Pho4 regulation. Since phosphatases produced by *C. albicans* as part of its low phosphate response are secreted, we hypothesized that their production would affect *P. aeruginosa* in co-culture, perhaps by providing access to phosphate freed from macromolecules and this model is discussed further below.

### In co-culture with ethanol-producing *C. albicans*, *P. aeruginosa* PhoB-plays independent roles in phosphate scavenging and phenazine-mediated antagonism

PhoB was important for *P. aeruginosa* growth in co-culture as Δ*phoB* formed fewer CFUs on *C. albicans* WT than *P. aeruginosa* WT (**Fig. 6E**). Notably, a comparable number of *P. aeruginosa* CFUs were recovered from *P. aeruginosa* WT and Δ*phz* suggesting the lack of phenazines was not a major reason for decreased CFU formation in Δ*phoB* but, likely, phosphate acquisition defects explained decreased growth in co-culture (**Fig. 6E**). The defect in CFU formation by Δ*phoB* compared to WT or Δ*phz* held true on *C. albicans* with a complemented copy of *ADH1.* On the *C. albicans adh1*Δ/Δ mutant, however, *P. aeruginosa* CFUs were similar for WT, Δ*phoB*, and Δ*phz* suggesting that in the absence of *C. albicans* Adh1 activity, *P. aeruginosa* PhoB, and its roles in phosphate acquisition or phenazine production, were not necessary for fitness. This finding is consistent with the model that elevated phosphatase production by the *C. albicans adh1*Δ/Δ (**Fig. B and D**) may eliminate the need for *P. aeruginosa* to produce phosphatases.

*C. albicans* CFUs were enumerated from the same co-cultures. Consistent with previous reports on the antifungal properties of the phenazine 5-MPCA [25, 27], *C. albicans* had lower CFUs upon co-culture with *P. aeruginosa* WT compared to when co-cultured with Δ*phoB* or Δ*phz* (**Fig. 6F**). In *C. albicans adh1*Δ/Δ co-cultures that did not support *P. aeruginosa* PhoB-dependent phenazine production (**Fig. 4**), *C. albicans* CFUs were not different between co-cultures with *P. aeruginosa* WT, Δ*phoB* or Δ*phz*.

### *P. aeruginosa* and *C. albicans* asynchronously activate their low phosphate responses dependent on *C. albicans* Adh1-mediated ethanol production

While *P. aeruginosa* had higher activity of its low phosphate responsive regulator PhoB in the presence of ethanol-producing *C. albicans* WT (**Fig. 3G** and **4A**), *C. albicans* had higher activity of its low phosphate responsive transcription factor Pho4 in *C. albicans adh1*Δ/Δ (**Fig. 6**). To explicitly examine the relationship between the *P. aeruginosa* and *C. albicans* low phosphate responses in co-culture we performed a cross-species correlation analysis on the co-culture gene expression data for a subset of *P. aeruginosa* PhoB-regulated genes (clique 3) and a subset of *C. albicans* Pho4-regulated genes (those also DEGs between *C. albicans* WT and *adh1*Δ/Δ shown in **Fig. 6A**). There was a striking pattern of inverse correlations between *P. aeruginosa* PhoB-regulated genes and *C. albicans* Pho4-regulated genes (**Fig. 7A**, upper triangle). In addition to being of high magnitudes, many same-species correlations (white background) as well as cross-species correlations (grey background) were statistically significant as indicated by circles (**Fig. 7A**, lower triangle). Of the two groups of strongly correlated genes, the one composed entirely of *P. aeruginosa* genes was more highly expressed in co-culture of *P. aeruginosa* and *C. albicans* WT than co-cultures of *P. aeruginosa* and *C. albicans adh1*Δ/Δ. Conversely, the group composed of mostly *C. albicans* genes and including the four *P. aeruginosa* phosphate transport-associated genes was more highly expressed in co-cultures of *P. aeruginosa* and *C. albicans adh1*Δ/Δ than in co-cultures of *P. aeruginosa* and *C. albicans* WT (**Fig. 7A**, right-hand barplot).

**Figure 7.**
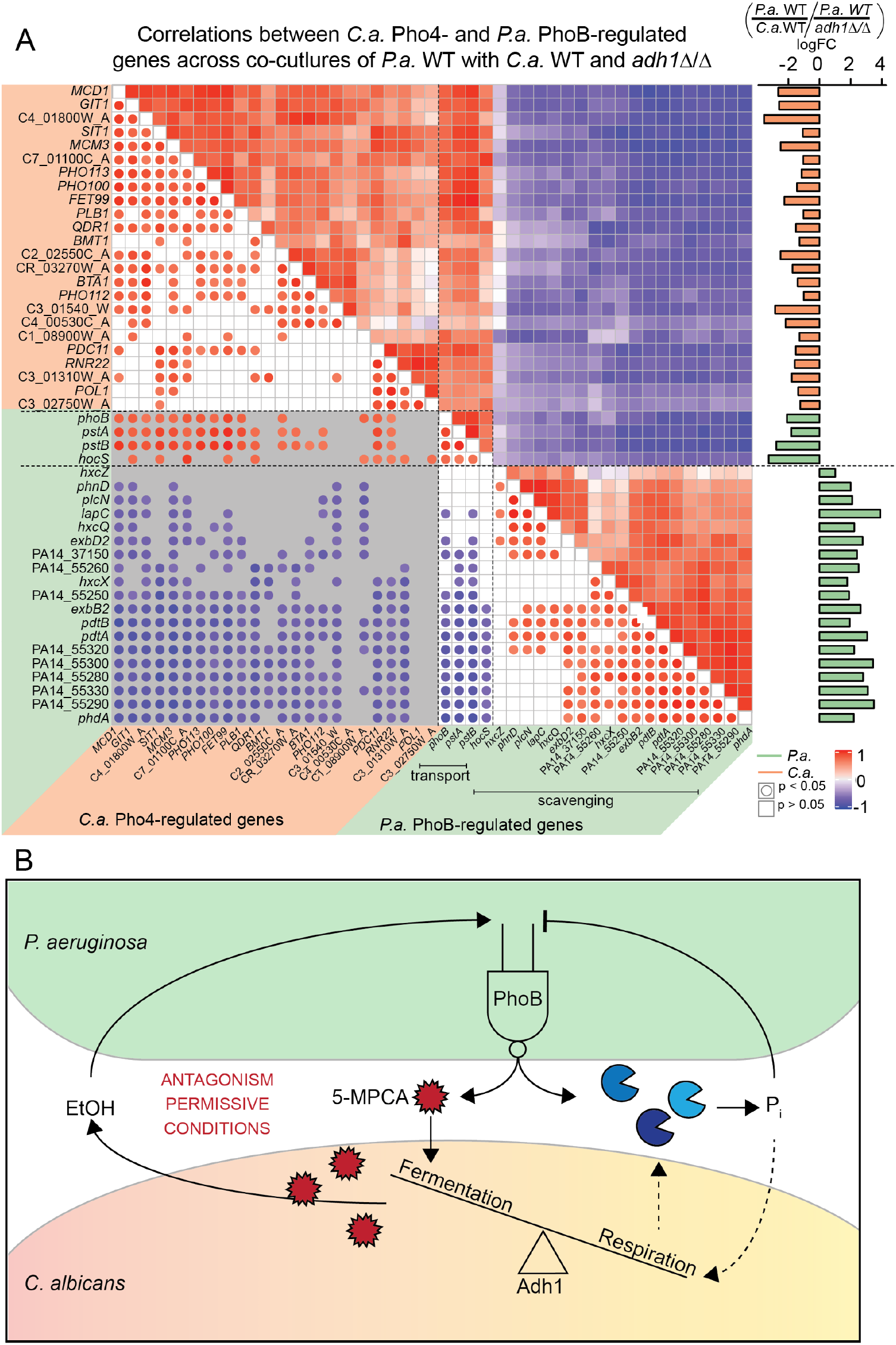
*P. aeruginosa* (*P.a.*) PhoB affects both *P.a.* and *C. albicans* (*C.a.*) fitness in co-culture through its control of phenazine production and phosphate acquisition. A) Pearson correlation analysis between *P.a.* (green annotations) and *C.a.* (orange annotations) low phosphate-responsive genes from co-cultures of *P.a.* WT with either *C.a.* WT or *adh1*Δ/Δ. Largely inverse relationships between *P.a.* PhoB- and *C.a.* Pho4-regulated genes is apparent. Log_2_FC (p<0.05) between *P.a*. with either *C.a*. WT or *C.a*. *adh1*Δ/Δ is shown in the right-hand bar plot. Lower half of correlogram shows which correlations are significant (filled circles) and indicates correlation values by color intensity relative to scale. Same species comparisons have white backgrounds and cross-species correlations have grey backgrounds. B) Model of PhoB activity in *P.a.-C.a.* co-cultures. PhoB mediates the conditional production of the antagonistic, antifungal phenazine 5-MPCA in response to phosphate and fungal ethanol production.**,p<0.01 by ANOVA.

The asynchronous and seemingly inverse activations of the *P. aeruginosa* and *C. albicans* low phosphate responses in co-culture, in combination with the differences in requirement for PhoB on *C. albicans* WT versus *adh1* (**Fig. 6E and 6F**), suggest that the two microbes were not responding to common environmental stimuli but rather that they influenced each other. The increased phosphatase activity in *C. albicans adh1*Δ/Δ even in the absence of *P. aeruginosa* (**Fig. 6D**) suggested that activation of the *C. albicans* low phosphate response may have inhibited activation of the *P. aeruginosa* low phosphate response providing negative regulation that, in concert with the lack of ethanol production, led to low PhoB activity in *P. aeruginosa* in co-culture with *C. albicans adh1*Δ/Δ and a consequent lack of 5-MPCA-derived red pigment accumulation (**Fig. 7B**).

## Discussion

### Co-culture transcriptomics explored dynamic microbe-microbe interactions

Whole genome transcriptional profiling has been used as a powerful tool to explore microbe-microbe interactions. By considering the gene expression patterns of both *P. aeruginosa* and *C. albicans* in co-culture, we found that the production of secreted phosphatases, a common good, was anti-correlated between *P. aeruginosa* and *C. albicans* depending on the ability of *C. albicans* to produce ethanol (**Fig 7A**). This interesting observation suggests that the two microbes interact differently in high and low phosphate environments and the dynamics of phosphate sensing, transport and enzymatic release by one organism can influence the microbial behavior of the other in complex ways that lead to the emergence of conditional antagonism.

Exploration of co-culture expression data sparked the investigation showing that signals of biologically-available phosphate and sub-inhibitory concentrations of ethanol were integrated into a PhoB-coordinated transcriptional response in *P. aeruginosa.* Interestingly, analysis of *C. albicans* gene expression in co-culture suggested these stimuli may be linked as well. Inorganic phosphate is well established as a negative regulator of the low phosphate response via the phosphate transport complex in *P. aeruginosa*, and here we show that ethanol, a common microbial fermentation product, positively regulated the *P. aeruginosa* low phosphate response specifically promoting the expression of secreted phosphatases, phospholipases, and antifungal phenazines. While these enzymes are only a subset of the PhoB regulon, PhoB activity and their expression were critical for P. aeruginosa in co-culture. In co-culture, this small armory of enzymes could make phosphate available by actively sequestering it away from neighboring microbes or by means of direct antagonism.

### eADAGE-based analysis identified novel transcriptional signals and increased interpretability of *P. aeruginosa* co-culture transcriptomics

The eADAGE model comprises 600 data-derived, weighted gene sets called signatures that are defined by the features of a hidden layer in an ensemble denoising autoencoder neural network (www.adage.greenelab.com). The signatures were learned from an unlabeled compendium of *P. aeruginosa* gene expression data and thus were defined solely on gene expression relationships rather than metadata. Fortuitously, many eADAGE signatures are enriched in one or more KEGG pathways [64] but others are not enriched in any. In this way, eADAGE provides an opportunity to expand the breadth of gene expression analysis beyond previously characterized pathways.

Despite our incomplete understanding of condition-dependent PhoB activity, we were able to identify a novel signal for a subset of PhoB-regulated genes through transcriptional profiling of *P. aeruginosa* grown in co-culture with *C. albicans* using eADAGE. Representation of the RNA-Seq data in reduced feature space yielded 48 differentially active signatures that distilled gene expression patterns amongst complex and dynamic systems. By analyzing eADAGE node activity, we were able to identify subtle signals for which genes did not individually meet the cutoffs for DEGs and whose pathways were not detected through over-representation analysis. Subtle changes in gene expression could still have dramatic phenotypic effects; for example, although the ethanol catabolism genes were not included in any enriched pathway, they were present in eADAGE differentially active signatures, and we identified ExaA-based ethanol catabolism as a potential regulator of the PhoB-mediated ethanol response including 5-MPCA production [91, 92]. Similarly, we observed enrichment of genes involved in isoprenoid metabolism which may be indicative of *P. aeruginosa* response to *C. albicans*-produced farnesol [18, 28, 93].

Of the 600 eADAGE signatures, only four contained over-representations of PhoB-controlled genes and while Node108neg was the only signature that reached the threshold for DASs, the activities of the other three trended upwards. The dynamics of co-culture that result in the constant utilization and solubilizing of phosphate differ from the steady state of phosphate limitation achievable in laboratory conditions. Therefore, while the low phosphate responses described in functional databases and the experimentally determined regulon did not show uniform increases in expression of the relevant genes, the data-defined gene signatures of eADAGE suggested that a tightly correlated sub-group of the Pho regulon is coordinately upregulated in response to ethanol-producing *C. albicans*.

It is only through the gene expression patterns observed by machine learning models trained on publicly available *P. aeruginosa* gene expression data that we were able to identify co-culture induced, ethanol-dependent PhoB activity. Considering the absence of samples from *P. aeruginosa* grown with *C. albicans* in the compendium of publicly available data, the distillation of the transcriptional pattern in Node108neg and its strong ethanol-induced activity in co-culture highlight the power of using unsupervised machine learning methods, in conjunction with a compendium of versatile conditions, to identify higher order gene expression patterns that are not evident in linear correlation based analyses and have not yet been manually annotated or systematically described.

### A conditional antagonism: PhoB-regulated antifungal production was dependent on *C. albicans* ethanol-production and phosphate limitation

Here we have shown that *P. aeruginosa* production of the antifungal phenazine 5-MPCA is dependent on PhoB, but canonical activation of PhoB by de-repression (Δ*pstB*) is insufficient for 5-MPCA production. We found that *P. aeruginosa* 5-MPCA production required ethanol, either produced by *C. albicans* or supplied exogenously, as an additional stimulus. While the mechanism by which ethanol influences PhoB activity is beyond the scope of this paper, PhoB has been implicated in cases of non-canonical signaling including auto-phosphorylation, promiscuity with non-canonical histidine kinases, activation in response to low iron, and stimulation by alternative phospho-donors such as acetyl-phosphate [35–39, 41, 42, 44, 55, 94–99]. The Pho regulon has been well characterized for direct PhoB targets [55] but also shown to have additional indirect targets across various low phosphate media [44]. Co-culture may be an environment conducive to alternative PhoB stimulation by ethanol. The dual requirement of both phosphate limitation and ethanol stimulation for *P. aeruginosa* 5-MPCA production presents a conditionally antagonistic relationship between *P. aeruginosa* and *C. albicans* where the degree to which the organisms cooperate, compete or target each other depends on metabolic cues and the abundance of an essential nutrient. Here we have presented a foundational example of an important and emerging paradigm that seeks to understand how environmental stimuli modulate microbial interactions.

## Materials and Methods

### Strains and growth conditions

Bacterial strains and plasmids used in this study are listed in **Table S3**. Bacteria were maintained on LB (lysogeny broth) with 1.5% agar [100]. Yeast strains were maintained on YPD (yeast peptone dextrose) with 2% agar. Where stated, ethanol (200-proof) was added to the medium (liquid or molten agar) to a final concentration of 1%. Planktonic cultures were grown on roller drums at 37°C for *P. aeruginosa* and at 30°C for C. *albicans*.

### Construction of in-frame deletions, complementation, and plasmids

Construction of plasmids, including in-frame deletion and complementation constructs, was completed using yeast cloning techniques in *Saccharomyces cerevisiae* as previously described [101] or Gibson assembly [102, 103]. Primers used for plasmid construction are listed in **Table S4**. In-frame deletion and single copy complementation constructs were made using the allelic replacement vector pMQ30 [101]. Promoter fusion constructs were made using a modified pMQ30 vector with *lacZ*-*GFP* fusion integrating at the neutral *att* site on the chromosome. The *pdtA* promoter region 199 bp upstream of the transcriptional start site (that included a PhoB binding site) was amplified from WT *P. aeruginosa* PA14 gDNA using the Phusion High-Fidelity DNA polymerase with primer tails homologous to the modified pMQ30 ATT KI vector containing tandem *lacZ-gfp* reporter genes. All plasmids were purified from yeast using Zymoprep™ Yeast Plasmid Miniprep II according to manufacturer’s protocol and transformed into electrocompetent *E. coli* strain S17/λpir by electroporation. Plasmids were introduced into *P. aeruginosa* by conjugation and recombinants were obtained using sucrose counter-selection. Genotypes were screened by PCR and confirmed by sequencing.

### Co-culture and mono-culture and colony biofilms

Co-cultures were inoculated first with 300 μl of a *C. albicans* culture in YPD grown for ∼16 then diluted in dH_2_O to OD_600_ = 5, which was bead spread onto YPD plates. *C. albican*s mono-cultures were inoculated with 5 μl of the same cell suspension as spots on YPD plates. *C. albicans* cultures were incubated 16 hours at 30°C then 24 hours at room temperature (∼23°C). Then, 5 μl of *P. aeruginosa* suspension, prepared from a 5 mL culture in LB grown for ∼16 then diluted in dH_2_O to OD_600_ = 2.5, was spotted on top of the *C. albicans* lawns for co-cultures or onto YPD or MOPS minimal medium for mono-cultures. *P. aeruginosa* cultures were incubated for 16 hours at 30°C. For gradient plates (described below) 500 μl of *C. albicans* suspension was used and *P. aeruginosa* was spotted across as 12 evenly spaced 5 μl spots. All images were taken on a Canon EOS Rebel T6i camera. For visualization of siderophores, pictures were taken under UV light.

### RNA collection

Total RNA was harvested from *P. aeruginosa* mono-cultures, *C. albicans* mono-cultures and *P. aeruginosa* – *C. albicans* co-cultures grown as described above. All samples were collected as cores from agar plates: cores were taken using a straw, cells were suspended by shaking agar plugs in 1 mL dH_2_0 on the disrupter genie for three minutes. Cells were spun down and resuspended in 1 mL Trizol and lysed by bead beating with mixed sizes of silicon beads on the Omni Bead Ruptor. Centrifugation induced phase separation and RNA was extracted from the aqueous phase where it was subsequently precipitated out with isopropanol and linear acrylamide. RNA was pelleted, washed with 70% ethanol and resuspended in nuclease-free dH_2_0 then stored at −80°C. Samples were prepared for sequencing with DNase treatment, ribodepletion and library preparation in accordance with Illumina protocols. Samples were barcoded and multiplexed in a NextSeq run for a total of 4.7×10^8^ reads.

### RNA-Seq Processing

Reads were processed using the CLC Genomics Workbench wherein reads were trimmed and filtered for quality using default parameters. For co-cultures, reads were first aligned to the *P. aeruginosa* UCPBB_PA14 genome from www.pseudomonas.com.

Then, all unaligned reads were aligned to *C. albicans* SC5314 genome Assembly 22 from www.candidagenome.org. For mono-cultures reads were only aligned to their appropriate reference. Results were exported from CLC including total counts, CPM and TPM. R was used for principle component (prcomp, stats library [104]) and consequent plotting (autoplot, ggplot2 [105]) of gene expression TPM data.

R was also used for differential gene expression analysis, EdgeR was used to process both *P. aeruginosa* and *C. albicans* gene expression separately [106]. Generalized linear models with mixed effect data design matrices were used to calculate fold-changes, p-values and FDRs for each comparison of interest. Volcano plots using EdgeR output (log_2_fold-change and -log_10_(FDR)) were made in R as well (ggplot2 [105]). Pathway over-representation analysis was carried out using *P. aeruginosa*- and *C. albicans*-associated KEGG pathways (ADAGEpath [64] and KEGGREST [107]) and calculated using a one-sided hypergeometric test (phyper, stats) with Bonferroni correction for multiple hypothesis testing (p.adjust, stats) [104].

### Accession Number

Data for our RNA-Seq analysis of *P. aeruginosa* and *C. albicans* gene expression in co-culture has been uploaded to the GEO repository (https://www.ncbi.nlm.nih.gov/geo/) with the accession number GSE148597.

### eADAGE analysis

We performed an eADAGE analysis in accordance with the workflow published in the R package [64]. Briefly, each gene expression profile in counts per million (CPM) from our RNA-Seq experiment was used to calculate a lower dimensional representation of the data called a signature activity profile. Then, differentially active signatures were identified by linear model. (limma, stats) This resulted in a set of signatures that were significantly different, but which may have been redundant. We applied pareto front optimization of minimal p-value and maximal absolute fold-change to arrive at a set of candidate signatures that exhibit statistically significant differences and less redundancy. Heatmaps show gene expression (CPM) or eADAGE signature activity scaled by feature (gene or signature) and are hierarchically clustered by the complete method using Euclidean distance by sample (ComplexHeatmap [108]).

### NanoString analysis

NanoString analysis was done using RNA isolated as above (without DNAse treatment) and 100 ng were applied to the codeset PaV5 (sequences for probesets used in this study in **Supplemental Dataset 4**) and processed as previously reported [2]. Counts were normalized to the geometric mean of spiked-in technical controls and five housekeeping genes (*ppiD*, *rpoD*, *soj*, *dnaN*, *pepP*, *dapF*). Normalized counts were used for heatmap construction and fold-change calculations.

### Measurement of β-galactosidase in reporter fusion strains

For co-culture promoter activity assays, *C. albicans* lawns were grown as described for RNA-Seq. *P. aeruginosa* was inoculated onto two filters placed on the *C. albicans* lawns to allow for interaction through diffusible compounds but separation of cells for quantification of promoter activity. *P. aeruginosa* cells were suspended in PBS by disrupter genie and diluted to OD_600_ = 0.05. β-Galactosidase (β-Gal) activity was measured as described by Miller [109]. β-Gal activity was measured in P. aeruginosa WT and normalized to that in Δ*phoB* which acted as a negative control.

### pNPP

For quantification of AP activity, we used a colorimetric assay using p-Nitrophenyl phosphate (pNPP) (NEB) as a substrate. Briefly, 5 μl of *P. aeruginosa* overnight culture were inoculated onto MOPS 0.7 mM phosphate agar plates with and without 1% ethanol and incubated at 37°C for 16 hours. Colony biofilms were collected from filters as described for the promoter fusion assays. 100 μl of cell suspension was added to 900 μl of 0.01 M Tris-HCl pH 8 buffer. After the addition of 25 μl 0.1% SDS and 50 μl chloroform cells incubated at 30 ° for 10 minutes. 30 μl of the aqueous phase was transferred to a 96 well plate containing 15 μl reaction buffer (5 μl 0.5 mM MgCl2 and 10 μl 1 M Tris pH 9.5) where 5 μl pNPP was added [36]. After 30 minutes OD_405_ was read on a plate reader (SpectraMax M2) and AP activity units were calculated as 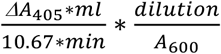 where 10.67 is the extinction coefficient, normally 18.5, adjusted to path length of the microtiter dish.

### Gradient plates

Phosphate gradient plates were made similarly to previously reported methods of creating pH gradient plates [110]. For YPD-based phosphate gradients plates used for co-cultures, first 32 mL of molten YPD+10mM potassium phosphate pH6 were poured into a 10 cm square petri dish (Corning, BP124-05) that rested in a custom 3D-printed prop that held the plate slanted at 30°. Once the bottom layer had solidified, the plate was laid flat and 32 mL of molten YPD without phosphate supplementation agar were poured atop. For MOPS-based gradient plates used for *P. aeruginosa* mono-cultures the procedure was the same except the first layer was 32 mL MOPS minimal media with 1 mM phosphate and the top layer was 32 mL MOPS minimal medium 0.4 mM phosphate. When needed, BCIP (Sigma-Aldrich #1158002001) (stock solution 60 mg/10 mL DMF) was added to a final molarity of 6 nM.

## Supporting information

Supp_dataset_1_Pa_on_CAF2vadh1_DEgenes

Supp_dataset_2_Ca_CAF2vadh1_DEgenes

Supp_dataset_3_Pa_and_Ca_CAF2vadh1_coculture_KEGGpathways

Supp_dataset_4_gene_lists

## Acknowledgements

Research reported in this publication was supported by grants from the National Institutes of Health to D.A.H. (R01 GM108492 to D.A.H) and HOGAN19G0, and NSF 1458359 (D.A.H. and G.D.). Support for G.D. came in part from 5T32GM008704 (G.D.). NIGMS P20GM113132 through the Molecular Interactions and Imaging Core (MIIC). STANTO19R0 from the Cystic Fibrosis Foundation and NIDDK P30-DK117469 (Dartmouth Cystic Fibrosis Research Center). RNA-Seq and NanoString were carried out at Dartmouth Medical School in the Genomics Shared Resource, which was established by equipment grants from the NIH and NSF and is supported in part by a Cancer Center Core Grant (P30CA023108) from the National Cancer Institute. We also thank Casey Greene and Alex Lee for a critical reading of the manuscript, Alan Collins for the gradient plate prop and Dianne Newman and Marian Llamas for generously sending strains.

## Supplemental Figure Legend

**Figure S1.**
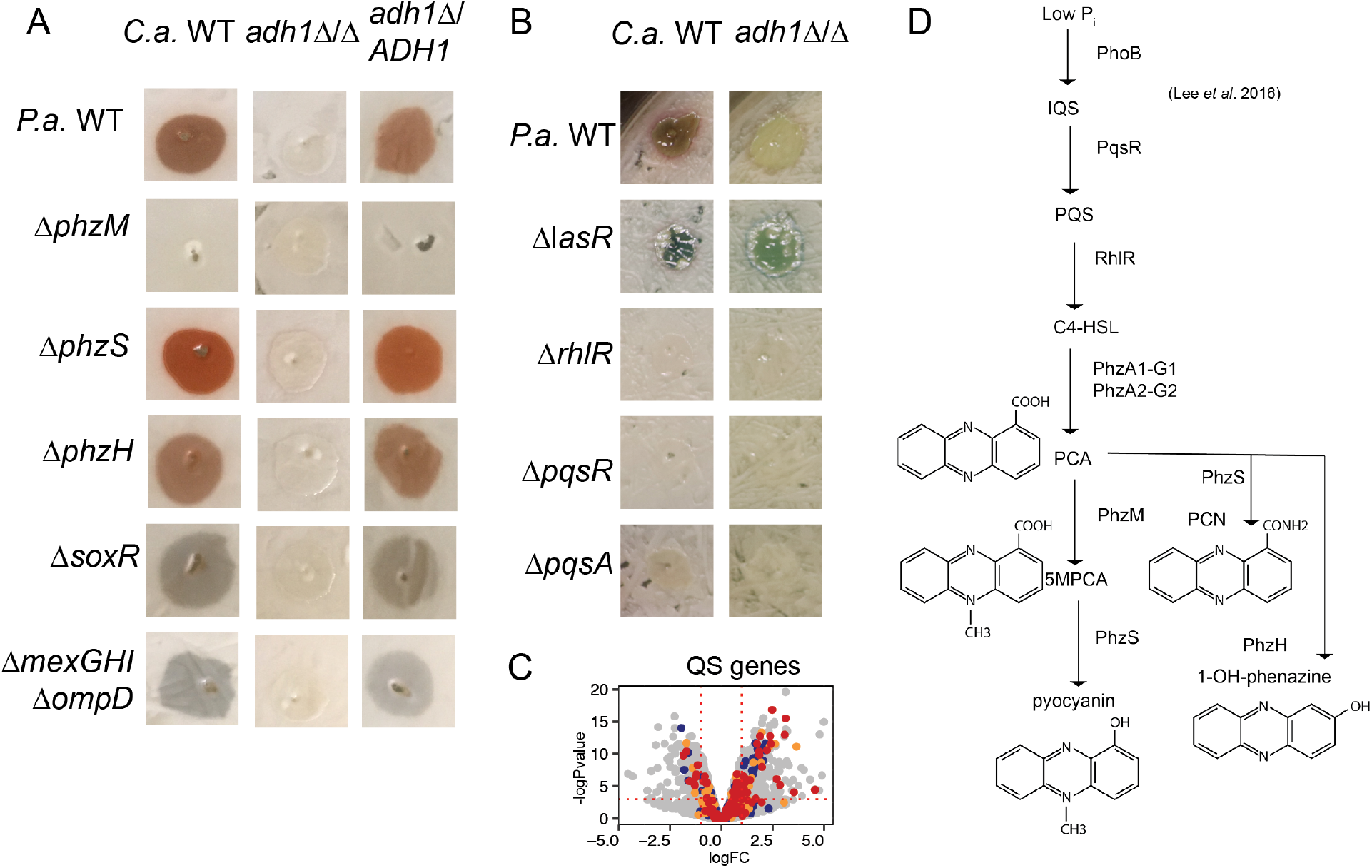
Red pigment formation is dependent on phenazine biosynthesis genes, phenazine transport genes and quorum sensing (QS) pathways in *P. aeruginosa* (*P.a.*). A) Co-cultures of *P.a*. wild type (WT) and mutants lacking genes involved in phenazine biosynthesis were inoculated onto lawns of *C.a.* (WT), *C.a. adh1*Δ/Δ and *adh1*Δ/Δ reconstituted with *ADH1* then incubated for 24 h. *P.a.* 5-MPCA phenazine biosynthesis (evident by red color) was not observed with the Δ*phzM* but was still produced by Δ*phzS* and Δ*phzH*. 5-MPCA production was dependent on the oxidative stress response gene *soxR* and 5-MPCA transport complex *mexGHI-ompD*. For all *P.a.* strains, 5-MPCA was only produced on *C.a.* with intact *ADH1*. B) Co-cultures of *P.a*. wild type (WT) and mutants lacking genes involved in quorum sensing were inoculated onto lawns of *C.a.* (WT), *C.a. adh1*Δ/Δ and *adh1*Δ/Δ reconstituted with *ADH1* then incubated for 48 h. *P.a.* mutants defective in QS pathways (Δ*lasR*, Δ*rhlR*, Δ*pqsR*, Δ*pqsA*) form less red pigment that *P.a.* WT on *C.a.* WT and no strains form red pigment on *C.a. adh1*Δ/Δ. C) Gene expression of *P.a*. LasR (blue), RhlR (orange) and PqsR (red) regulated genes had heterogenous expression with genes both up and down regulated. D) Red pigment formation being dependent on QS pathways of RhlR and PqsR is consistent with integration of QS with PhoB via the integrative quorum sensing (IQS) pathway in which low phosphate triggers PhoB activity which influences PqsR and RhlR to act, via their cognate autoinducers PQS and C4-HSL respectively, in a regulatory cascade eventually promoting the transcription of phenazine biosynthesis genes (*phzA1-G1*, *phzA2-G2*) and consequent phenazine carboxylic acid (PCA) production. PhoB and QS have also been reported to effect the expression of phenazine modification genes *phzM, phzS,* and *phzH* necessary for the conversion of PCA to 5-methyl-phenazine-1-carboxylic acid (5-MPCA), pyocyanin, phenazine-1-carboxamide (PCN) and 1-hydroxy-phenazine (1-OH-phenazine) [74].

## Supporting Information

**S1 Table.** Clustering coefficients for co-culture data.

| Groups | CC for gene expression | SD | CC for signature activity | SD |
| --- | --- | --- | --- | --- |
| CAF2 vs *adh1Δ/Δ* | 0.38 |  | 0.68 |  |
| Random (5v5) | 0.14 | ± 0.07 | 0.12 | ± 0.12 |
| Coculture vs Monoculture | 0.42 |  | 0.53 |  |
| Random(10v3) | 0.14 | ± 0.16 | 0.15 | ± 0.22 |
| CAF2 vs YPD | 0.58 |  | 0.80 |  |
| Random (5v3) | 0.13 | ± 0.09 | 0.10 | ± 0.11 |

**S2 Table.** eADAGE gene-gene network cliques of DEGs from co-culture.

| Clique Number | Description | Clique size | Genes |
| --- | --- | --- | --- |
| 1 | Phenazine biosynthesis | 17 | PA0522, PA3039, PA5383, PA1219, PA1221, PA1220, PA1216, PA1218, *phzA1*, *phzG2*, *phzE2*, *phzC2*, *phzD2*, *phzM*, *phzS*, *phzB1*, *phzF2* |
| 2 | Pyochelin biosynthesis | 29 | *fepD*, *fepB*, PA4155, *foxl*, *pvdS*, *femI*, *fiuI*, *femR*, PA5217, PA1301 PA1300, *hasAP*, PA4570, *pchR*, *fumC1*, *sodM*, PA4471, *fiuR*, *pchA*, *ampO*, *pchD*, *pchB*, *fptA*, *pchC*, *pchE*, PA4220, PA4222, PA4223, *pchF* |
| 3 | Phosphate transport and acquisition | 23 | PA2120, *hocS*, *phoB*, *pstA*, *pstC*, *plcN*, PA3383, *exbB2*, *phdA*, *pdtB*, PA0701, PA0699, *hxcX*, *lapA*, *exbD2*, PA0696, PA0697, PA0698, *hxcQ*, PA0700, *hxcZ*, PA0695, *pdtA* |
| 4 | Isoprenoid metabolism | 7 | *liuD*, *liuB*, *mmsB*, PA2557, PA2553, *liuC*, *liuA* |
| 5 | Magnesium transport | 6 | *mgtA*, PA4826, PA4635, PA4822, PA4824, PA4823 |
| 6 | Aconitate porin | 4 | *opdH*, PA0753, PA0752, PA0754 |
| 7 | Pyrimidine metabolism | 3 | *dht*, PA0440, PA0439 |
| 8 | Pyocin Biosynthesis | 3 | PA0629, PA0630, PA0637 |
| 9 | Spermidine biosynthesis | 3 | PA4773, PA4774, PA4782 |
| 10 | Histidine catabolism | 2 | *hutH*, PA5096 |
| 11 | Heat shock response | 2 | *hscA*, *fdx2* |
| 12 | Reactive oxygen stress response | 2 | *ahpF*, *katB* |

**S3 Table.** Strains and plasmids used in this study.

| Strain | Lab stock# | Strain description | Source |
| --- | --- | --- | --- |
| ***P. aeruginosa*** |  |  |  |
| PA14 WT | DH123 | Laboratory reference strain | [1] |
| PA14 $\Delta phoB$ | DH3599 | Deletion mutant in *phoB* | This study |
| PA14 $\Delta phoB + phoB$ | DH3600 | Native locus complementation of *phoB* | This study |
| PA14 $\Delta phoR$ | DH3774 | Deletion mutant of *phoR* | [2] |
| PA14 $\Delta phoB + phoR$ | DH3755 | Plasmid-based arabinose inducible over-expression vector of *phoR* | This study |
| PA14 $\Delta pstB$ | DH3601 | Deletion mutant of *pstB* | This study |
| PA14 *pstB::TnM* | DH753 | Transposon insertion mutant of *pstB* | [3] |
| PA14 $\Delta phz$ | DH933 | Deletion mutant of *phzA1-G1* and *phzA2-G2* | [4] |
| PA14 $\Delta phzA1$ | DH1728 | Deletion mutant of *phzA1* | (Dietrich, Price-Whelan et al. 2006) |
| PA14 $\Delta phzA2$ | DH1735 | Deletion mutant of *phzA2* | (Dietrich, Price-Whelan et al. 2006) |
| PA14 $\Delta phzM$ | DH944 | Deletion mutant of *phzM* | [5] |
| PA14 $\Delta mexGH1\Delta ompD$ | DH1376 | Deletion mutant of *mexGH1* and *ompD* | [5] |
| PA14 $\Delta soxR$ | DH1377 | Deletion mutant of *soxR* | [5] |
| PA14 $\Delta exaA$ | DH2256 | Deletion mutant of *exaA* | [6] |
| PA14 $\Delta exaA + exaA$ | DH2677 | Native locus complementation of *exaA* | [6] |
| PA14 WT PpdtA::lacZ-gfp | DH3780 | Promoter fusion reported construct of PpdtA chromosomally integrated at the *att* site | This study |
| PA14 $\Delta phoB$ PpdtA::lacZ-gfp | DH3781 | Promoter fusion reported construct of PpdtA chromosomally integrated at the *att* site in $\Delta phoB$ background | This study |
| CAF2 WT | DH48 | Laboratory reference strain | [7] |
| CAF2 *adh1*$\Delta/\Delta$ | DH2236 | Homozygous deletion mutant of *ADH1* | [8] |
| CAF2 *adh1*$\Delta/\Delta$ + *ADH1* | DH2177 | Heterozygous native locus single allele complementation of *ADH1* | [8] |
| PA14 $\Delta ackA$ | DH3782 | Deletion mutant of *ackA* | [9] |
| PA14 $\Delta ackA \Delta pta$ | DH3783 | Deletion mutant of *ackA* and *pta* | [9] |
| PA14 $\Delta ackA \Delta pta \Delta phz$ | Dh3784 | Deletion mutant of *ackA*, *pta*, *phzA1-G1* and *phzA2-G2* | [9] |
| PA14 *exaA::TnM* | DH2130 | ethanol catabolic transposon insertion mutant | [10] |
| PA14 *pqqB::TnM* | DH2131 | ethanol catabolic transposon insertion mutant | [10] |
| PA14 *acsA::TnM* | DH2132 | ethanol catabolic transposon insertion mutant | [10] |
| PA14 $\Delta kinB$ | DH3778 | Deletion mutant of *kinB* | [2] |
| PA14 $\Delta kinB$ + *kinB* | DH3779 | Plasmid-based arabinose inducible over-expression vector of *kinB* | This study |
| PAO1 | DH3283 | Laboratory reference strain | [11] |
| PAO1 $\Delta phoB$ | DH3284 | Deletion mutant of *phoB* | [11] |
| PAO1 $\Delta vreA$ | DH3285 | Deletion mutant of *vreA* | [11] |
| PAO1 $\Delta vreI$ | DH3286 | Deletion mutant of *vreI* | [11] |
| PAO1 $\Delta vreR$ | DH3287 | Deletion mutant of *vreR* | [11] |
| PAO1 $\Delta phoB \Delta vreR$ | DH3288 | Deletion mutant of *phoB* and *vreR* | [11] |
| ***E. coli*** |  |  |  |
| S17 λpirS | DH71 | Mating competent strain used for plasmid conjugation | |
| **Plasmids** |  |  |  |
| pMQ30 |  | allelic replacement vector, GmR | [12] |
| *phoB* complement |  |  | This study |
| pMQ72 |  | Arabinose inducible over-expression plasmid, empty vector | [12] |
| *phoR* OE |  | Arabinose inducible over-expression plasmid, containing *phoR* | This study |
| pHERD20 |  | Arabinose inducible over-expression plasmid | [13] |
| kinB OE |  | Arabinose inducible over-expression plasmid, containing *kinB* | [14] |
| *PpdtA-lacZ* |  | *PpdtA-lacZ* promoter fusion, GmR in S17 *E. coli* | This study |
| pEX18-Gm |  | Suicide vector for allelic replacement, GmR | [15] |

**S4 Table.**
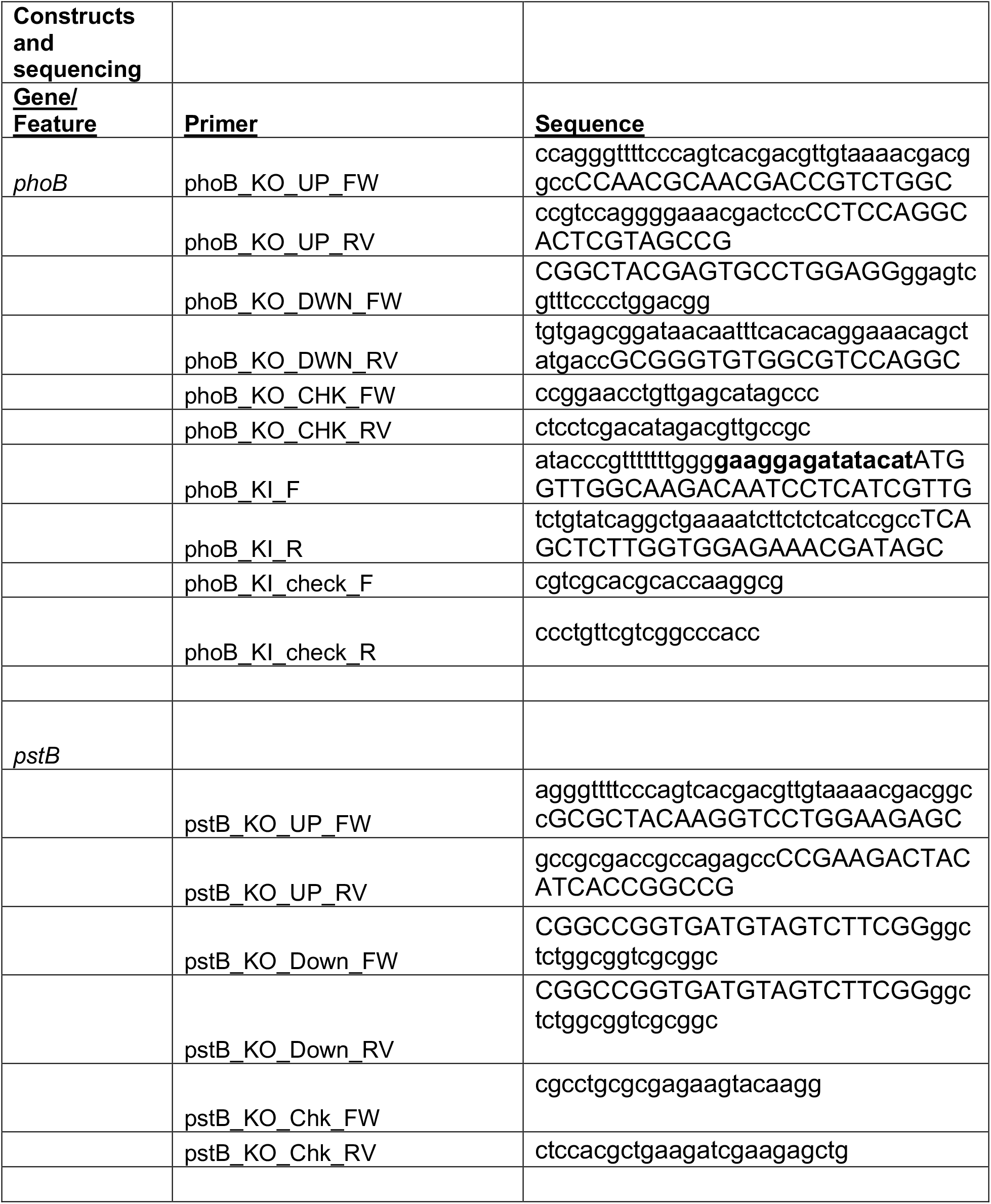
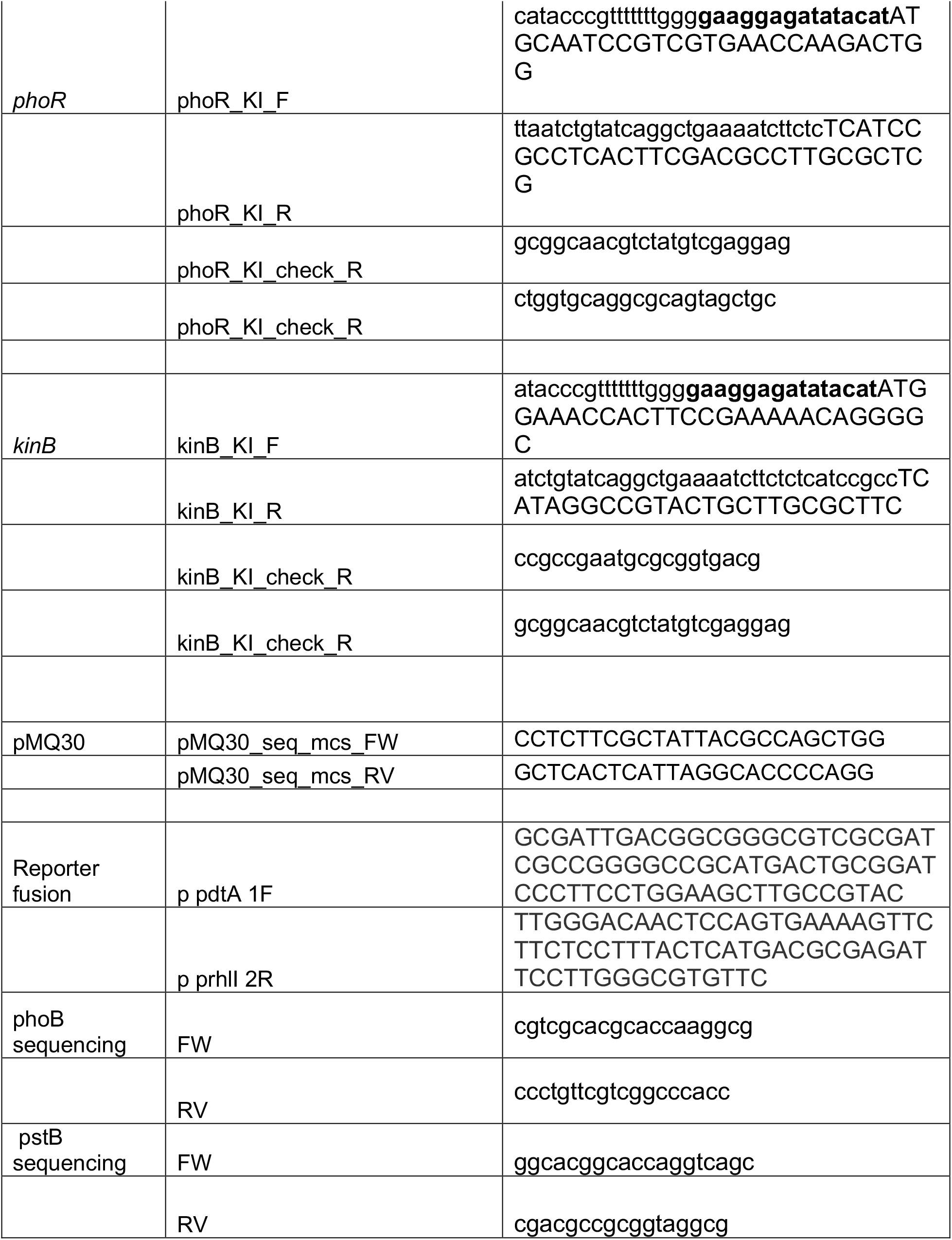
Primers used in this study.

**S1 Dataset. P.a. DEGs in co-culture with C.a..**

**S2 Dataset. C.a. DEGS in co-culture with P.a..**

**S3 Dataset. KEGG pathway analyses.**

**S4 Dataset. Gene sets used throughout the paper.**

## References

1. Hughes WT, Kim HK. Mycoflora in cystic fibrosis: some ecologic aspects of *Pseudomonas aeruginosa* and *Candida albicans*. Mycopathol Mycol Appl. 1973;50(3):261–9. Epub 1973/07/31. PubMed PMID: 4199669.

2. Grahl N, Dolben EL, Filkins LM, Crocker AW, Willger SD, Morrison HG, et al. Profiling of bacterial and fungal microbial communities in cystic fibrosis sputum using RNA. mSphere. 2018;3(4):e00292–18. doi: 10.1128/mSphere.00292-18. PubMed PMID: 30089648.

3. Azoulay E, Timsit J-F, Tafflet M, de Lassence A, Darmon M, Zahar J-R, et al. *Candida c*olonization of the respiratory tract and subsequent *Pseudomonas* ventilator-associated pneumonia. Chest. 2006;129(1):110–7. doi: https://doi.org/10.1378/chest.129.1.110.

4. Falleiros de Padua RA, Norman Negri MF, Svidzinski AE, Nakamura CV, Svidzinski TI. Adherence of *Pseudomonas aeruginosa* and *Candida albicans* to urinary catheters. Rev Iberoam Micol. 2008;25(3):173–5. Epub 2008/09/13. doi: 10.1016/s1130-1406(08)70040-8. PubMed PMID: 18785788.

5. Gupta N, Haque A, Mukhopadhyay G, Narayan RP, Prasad R. Interactions between bacteria and *Candida* in the burn wound. Burns. 2005;31(3):375–8. doi: https://doi.org/10.1016/j.burns.2004.11.012.

6. Nseir S, Jozefowicz E, Cavestri B, Sendid B, Di Pompeo C, Dewavrin F, et al. Impact of antifungal treatment on *Candida*–*Pseudomonas* interaction: a preliminary retrospective case–control study. Intensive Care Med. 2007;33(1):137–42. doi: 10.1007/s00134-006-0422-0.

7. Pierce GE. Pseudomonas aeruginosa, Candida albicans, and device-related nosocomial infections: implications, trends, and potential approaches for control. J Ind Microbiol Biotechnol. 2005;32(7):309–18. doi: 10.1007/s10295-005-0225-2.

8. Kerr JR. Suppression of fungal growth exhibited by *Pseudomonas aeruginosa*. J Clin Microbiol. 1994;32(2):525–7. PubMed PMID: 8150966.

9. Bakare N, Rickerts V, Bargon J, Just-Nubling G. Prevalence of *Aspergillus fumigatus* and other fungal species in the sputum of adult patients with cystic fibrosis. Mycoses. 2003;46(1-2):19–23. Epub 2003/02/18. doi: 10.1046/j.1439-0507.2003.00830.x. PubMed PMID: 12588478.

10. Kaleli I, Cevahir N, Demir M, Yildirim U, Sahin R. Anticandidal activity of *Pseudomonas aeruginosa* strains isolated from clinical specimens. Mycoses. 2007;50(1):74–8. Epub 2007/02/17. doi: 10.1111/j.1439-0507.2006.01322.x. PubMed PMID: 17302753.

11. Bauernfeind A, Bertele RM, Harms K, Horl G, Jungwirth R, Petermuller C, et al. Qualitative and quantitative microbiological analysis of sputa of 102 patients with cystic fibrosis. Infection. 1987;15(4):270–7. Epub 1987/07/01. doi: 10.1007/bf01644137. PubMed PMID: 3117700.

12. Bandara H, Yau JYY, Watt RM, Jin LJ, Samaranayake LP. *Pseudomonas aeruginosa* inhibits *in-vitro Candida* biofilm development. BMC Microbiol. 2010;10(1):125. doi: 10.1186/1471-2180-10-125.

13. Bergeron AC, Seman BG, Hammond JH, Archambault LS, Hogan DA, Wheeler RT. *Candida albicans* and *Pseudomonas aeruginosa* interact to enhance virulence of mucosal infection in transparent zebrafish. Infect Immun. 2017;85(11):e00475–17. doi: 10.1128/iai.00475-17.

14. Brand A, Barnes JD, Mackenzie KS, Odds FC, Gow NAR. Cell wall glycans and soluble factors determine the interactions between the hyphae of *Candida albicans* and *Pseudomonas aeruginosa*. FEMS Microbiol Lett. 2008;287(1):48–55. doi: 10.1111/j.1574-6968.2008.01301.x.

15. Lopez-Medina E, Fan D, Coughlin LA, Ho EX, Lamont IL, Reimmann C, et al. *Candida albicans* Inhibits *Pseudomonas aeruginosa* Virulence through Suppression of Pyochelin and Pyoverdine Biosynthesis. PLoS Path. 2015;11(8):e1005129-e. doi: 10.1371/journal.ppat.1005129. PubMed PMID: 26313907.

16. Purschke FG, Hiller E, Trick I, Rupp S. Flexible survival strategies of *Pseudomonas aeruginosa* in biofilms result in increased fitness compared with *Candida albicans*. Molecular & Cellular Proteomics. 2012;11(12):1652–69. doi: 10.1074/mcp.M112.017673.

17. Trejo-Hernández A, Andrade-Domínguez A, Hernández M, Encarnación S. Interspecies competition triggers virulence and mutability in *Candida albicans*– *Pseudomonas aeruginosa* mixed biofilms. The ISME Journal. 2014;8(10):1974–88. doi: 10.1038/ismej.2014.53.

18. Cugini C, Morales DK, Hogan DA. *Candida albicans*-produced farnesol stimulates *Pseudomonas* quinolone signal production in LasR-defective *Pseudomonas aeruginosa* strains. Microbiology. 2010;156(Pt 10):3096–107. doi: 10.1099/mic.0.037911-0.

19. De Sordi L, Mühlschlegel FA. Quorum sensing and fungal–bacterial interactions in *Candida albicans*: a communicative network regulating microbial coexistence and virulence. FEMS Yeast Res. 2009;9(7):990–9. doi: 10.1111/j.1567-1364.2009.00573.x.

20. Fourie R, Ells R, Swart CW, Sebolai OM, Albertyn J, Pohl CH. *Candida albicans* and *Pseudomonas aeruginosa* Interaction, with Focus on the Role of Eicosanoids. Front Physiol. 2016;7(64). doi: 10.3389/fphys.2016.00064.

21. Hogan DA, Vik Å, Kolter R. A *Pseudomonas aeruginosa* quorum-sensing molecule influences *Candida albicans* morphology. Mol Microbiol. 2004;54(5):1212–23. doi: 10.1111/j.1365-2958.2004.04349.x.

22. Holcombe LJ, McAlester G, Munro CA, Enjalbert B, Brown AJP, Gow NAR, et al. *Pseudomonas aeruginosa* secreted factors impair biofilm development in *Candida albicans*. Microbiology. 2010;156(5):1476–86. doi: https://doi.org/10.1099/mic.0.037549-0.

23. McAlester G, O’Gara F, Morrissey JP. Signal-mediated interactions between *Pseudomonas aeruginosa* and *Candida albicans*. J Med Microbiol. 2008;57(Pt 5):563–9. Epub 2008/04/26. doi: 10.1099/jmm.0.47705-0. PubMed PMID: 18436588.

24. Sakhtah H, Koyama L, Zhang Y, Morales DK, Fields BL, Price-Whelan A, et al. The *Pseudomonas aeruginosa* efflux pump MexGHI-OpmD transports a natural phenazine that controls gene expression and biofilm development. Proc Natl Acad Sci U S A. 2016;113(25):E3538–47. doi: 10.1073/pnas.1600424113.

25. Morales DK, Jacobs NJ, Rajamani S, Krishnamurthy M, Cubillos-Ruiz JR, Hogan DA. Antifungal mechanisms by which a novel *Pseudomonas aeruginosa* phenazine toxin kills *Candida albicans* in biofilms. Mol Microbiol. 2010;78(6):1379–92. doi: 10.1111/j.1365-2958.2010.07414.x.

26. Morales DK, Grahl N, Okegbe C, Dietrich LEP, Jacobs NJ, Hogan DA. Control of *Candida albicans* metabolism and biofilm formation by *Pseudomonas aeruginosa* phenazines. mBio. 2013;4(1):e00526–12. doi: 10.1128/mBio.00526-12.

27. Gibson J, Sood A, Hogan DA. *Pseudomonas aeruginosa*-*Candida albicans* interactions: localization and fungal toxicity of a phenazine derivative. Appl Environ Microbiol. 2009;75(2):504–13. doi: 10.1128/AEM.01037-08.

28. Cugini C, Calfee MW, Farrow JM, Morales DK, Pesci EC, Hogan DA. Farnesol, a common sesquiterpene, inhibits PQS production in *Pseudomonas aeruginosa*. Mol Microbiol. 2007;65(4):896–906. doi: 10.1111/j.1365-2958.2007.05840.x.

29. Chen AI, Dolben EF, Okegbe C, Harty CE, Golub Y, Thao S, et al. *Candida albicans* ethanol stimulates *Pseudomonas aeruginosa* WspR-controlled biofilm formation as part of a cyclic relationship involving phenazines. PLoS Path. 2014;10(10):e1004480-e. doi: 10.1371/journal.ppat.1004480.

30. Kerr JR, Taylor GW, Rutman A, Høiby N, Cole PJ, Wilson R. *Pseudomonas aeruginosa* pyocyanin and 1-hydroxyphenazine inhibit fungal growth. J Clin Pathol. 1999;52(5):385–7. doi: 10.1136/jcp.52.5.385. PubMed PMID: 10560362.

31. Harty CE, Martins D, Doing G, Mould DL, Clay ME, Occhipinti P, et al. Ethanol stimulates trehalose production through a SpoT-DksA-AlgU-dependent pathway in *Pseudomonas aeruginosa*. J Bacteriol. 2019;201(12):e00794–18. doi: 10.1128/JB.00794-18.

32. Lewis KA, Baker AE, Chen AI, Harty CE, Kuchma SL, O’Toole GA, et al. Ethanol decreases *Pseudomonas aeruginosa* flagellar motility through the regulation of flagellar stators. J Bacteriol. 2019;201(18):e00285–19. Epub 2019/05/22. doi: 10.1128/jb.00285-19. PubMed PMID: 31109994.

33. DeVault JD, Kimbara K, Chakrabarty AM. Pulmonary dehydration and infection in cystic fibrosis: evidence that ethanol activates alginate gene expression and induction of mucoidy in *Pseudomonas aeruginosa*. Mol Microbiol. 1990;4(5):737–45. doi: 10.1111/j.1365-2958.1990.tb00644.x.

34. Aendekerk S, Diggle SP, Song Z, Hoiby N, Cornelis P, Williams P, et al. The MexGHI-OpmD multidrug efflux pump controls growth, antibiotic susceptibility and virulence in *Pseudomonas aeruginosa* via 4-quinolone-dependent cell-to-cell communication. Microbiology. 2005;151(Pt 4):1113–25. Epub 2005/04/09. doi: 10.1099/mic.0.27631-0. PubMed PMID: 15817779.

35. Bains M, Fernández L, Hancock REW. Phosphate starvation promotes swarming motility and cytotoxicity of *Pseudomonas aeruginosa*. Appl Environ Microbiol. 2012;78(18):6762–8. doi: 10.1128/AEM.01015-12.

36. Blus-Kadosh I, Zilka A, Yerushalmi G, Banin E. The effect of *pstS* and phoB on quorum sensing and swarming motility in *Pseudomonas aeruginosa*. PLoS One. 2013;8(9):e74444-e. doi: 10.1371/journal.pone.0074444.

37. Faure LM, Llamas MA, Bastiaansen KC, de Bentzmann S, Bigot S. Phosphate starvation relayed by PhoB activates the expression of the *Pseudomonas aeruginosa* vreI ECF factor and its target genes. Microbiology. 2013;159(Pt_7):1315–27. doi: 10.1099/mic.0.067645-0.

38. Haddad A, Jensen V, Becker T, HÃ¤ussler S. The Pho regulon influences biofilm formation and type three secretion in *Pseudomonas aeruginosa*. Environ Microbiol Rep. 2009;1(6):488–94. doi: 10.1111/j.1758-2229.2009.00049.x.

39. Jensen V, Löns D, Zaoui C, Bredenbruch F, Meissner A, Dieterich G, et al. RhlR expression in *Pseudomonas aeruginosa* is modulated by the *Pseudomonas* quinolone signal via PhoB-dependent and -independent pathways. J Bacteriol. 2006;188(24):8601–6. doi: 10.1128/JB.01378-06.

40. Lamarche MG, Wanner BL, Crépin S, Harel J. The phosphate regulon and bacterial virulence: a regulatory network connecting phosphate homeostasis and pathogenesis. FEMS Microbiol Rev. 2008;32(3):461–73. doi: 10.1111/j.1574-6976.2008.00101.x.

41. Quesada JM, Otero-Asman JR, Bastiaansen KC, Civantos C, Llamas MA. The activity of the *Pseudomonas aeruginosa* virulence regulator σVreI is modulated by the anti-σ factor VreR and the transcription factor PhoB. Front Microbiol. 2016;7:1159-. doi: 10.3389/fmicb.2016.01159.

42. Shoriridge VD, Lazdunski A, Vasil ML. Osmoprotectants and phosphate regulate expression of phospholipase C in *Pseudomonas aeruginosa*. Mol Microbiol. 1992;6(7):863–71. doi: 10.1111/j.1365-2958.1992.tb01537.x.

43. Zaborin A, Gerdes S, Holbrook C, Liu DC, Zaborina OY, Alverdy JC. *Pseudomonas aeruginosa* overrides the virulence inducing effect of opioids when it senses an abundance of phosphate. PLoS One. 2012;7(4):e34883-e. doi: 10.1371/journal.pone.0034883.

44. Zaborin A, Romanowski K, Gerdes S, Holbrook C, Lepine F, Long J, et al. Red death in *Caenorhabditis elegans* caused by *Pseudomonas aeruginosa* PAO1. Proc Natl Acad Sci U S A. 2009;106(15):6327–32. doi: 10.1073/pnas.0813199106.

45. Chand NS, Lee JS-W, Clatworthy AE, Golas AJ, Smith RS, Hung DT. The sensor kinase KinB regulates virulence in acute *Pseudomonas aeruginosa* infection. J Bacteriol. 2011;193(12):2989–99. doi: 10.1128/JB.01546-10.

46. Cornforth DM, Dees JL, Ibberson CB, Huse HK, Mathiesen IH, Kirketerp-Møller K, et al. *Pseudomonas aeruginosa* transcriptome during human infection. Proc Natl Acad Sci U S A. 2018;115(22):E5125–E34. doi: 10.1073/pnas.1717525115.

47. Cox CD, Adams P. Siderophore activity of pyoverdin for *Pseudomonas aeruginosa*. Infect Immun. 1985;48(1):130.

48. Damron FH, Oglesby-Sherrouse AG, Wilks A, Barbier M. Dual-seq transcriptomics reveals the battle for iron during *Pseudomonas aeruginosa* acute murine pneumonia. Sci Rep. 2016;6(1):39172-. doi: 10.1038/srep39172.

49. Damron FH, Qiu D, Yu HD. The *Pseudomonas aeruginosa* sensor kinase KinB negatively controls alginate production through AlgW-dependent MucA proteolysis. J Bacteriol. 2009;191(7):2285–95. Epub 2009/01/23. doi: 10.1128/JB.01490-08. PubMed PMID: 19168621.

50. Francis VI, Stevenson EC, Porter SL. Two-component systems required for virulence in *Pseudomonas aeruginosa*. FEMS Microbiol Lett. 2017;364(11). doi: 10.1093/femsle/fnx104.

51. Liu PV, Shokrani F. Biological activities of pyochelins: iron-chelating agents of *Pseudomonas aeruginosa*. Infect Immun. 1978;22(3):878–90. Epub 1978/12/01. PubMed PMID: 103839; PubMed Central PMCID: PMCPMC422240.

52. Schmidberger A, Henkel M, Hausmann R, Schwartz T. Influence of ferric iron on gene expression and rhamnolipid synthesis during batch cultivation of *Pseudomonas aeruginosa* PAO1. Appl Microbiol Biotechnol. 2014;98(15):6725–37. Epub 2014/04/23. doi: 10.1007/s00253-014-5747-y. PubMed PMID: 24752844.

53. Tan J, Doing G, Lewis KA, Price CE, Chen KM, Cady KC, et al. Unsupervised extraction of stable expression signatures from public compendia with an ensemble of neural networks. Cell Systems. 2017;5(1):63–71.e6. doi: 10.1016/J.CELS.2017.06.003.

54. Hogan DA, Kolter R. *Pseudomonas-Candida* interactions: an ecological role for virulence factors. Science (New York, NY). 2002;296(5576):2229–32. doi: 10.1126/science.1070784.

55. Bielecki P, Jensen V, Schulze W, Gödeke J, Strehmel J, Eckweiler D, et al. Cross talk between the response regulators PhoB and TctD allows for the integration of diverse environmental signals in *Pseudomonas aeruginosa*. Nucleic Acids Res. 2015;43(13):6413–25. doi: 10.1093/nar/gkv599.

56. Ching T, Himmelstein DS, Beaulieu-Jones BK, Kalinin AA, Do BT, Way GP, et al. Opportunities and obstacles for deep learning in biology and medicine. Journal of The Royal Society Interface. 2018;15(141):20170387. doi: doi:10.1098/rsif.2017.0387.

57. Greene CS, Foster JA, Stanton BA, Hogan DA, Bromberg Y. Computational approaches to study micorbes and microbiomes. Pac Symp Biocomput. 2016;21:557–67. doi: 10.1142/9789814749411_0051.

58. Tan J, Hammond JH, Hogan DA, Greene CS. ADAGE-based integration of publicly svailable *Pseudomonas aeruginosa* gene expression data with denoising autoencoders illuminates microbe-host interactions. mSystems. 2016;1(1):e00025–15. doi: 10.1128/mSystems.00025-15.

59. Taroni JN, Greene CS, Martyanov V, Wood TA, Christmann RB, Farber HW, et al. A novel multi-network approach reveals tissue-specific cellular modulators of fibrosis in systemic sclerosis. Genome Med. 2017;9(1):27. doi: 10.1186/s13073-017-0417-1.

60. Way GP, Greene CS. Extracting a biologically relevant latent space from cancer transcriptomes with variational autoencoders. Pac Symp Biocomput. 2018;23:80–91. doi: doi:10.1142/9789813235533_0008 10.1142/9789813235533_0008.

61. Zhu Q, Wong AK, Krishnan A, Aure MR, Tadych A, Zhang R, et al. Targeted exploration and analysis of large cross-platform human transcriptomic compendia. Nat Methods. 2015;12(3):211–4. doi: 10.1038/nmeth.3249.

62. Taroni JN, Grayson PC, Hu Q, Eddy S, Kretzler M, Merkel PA, et al. MultiPLIER: A transfer learning framework for transcriptomics reveals systemic features of rare disease. Cell Systems. 2019;8(5):380–94.e4. Epub 2019/05/24. doi: 10.1016/j.cels.2019.04.003. PubMed PMID: 31121115; PubMed Central PMCID: PMCPMC6538307.

63. Chen KM, Tan J, Way GP, Doing G, Hogan DA, Greene CS. PathCORE-T: identifying and visualizing globally co-occurring pathways in large transcriptomic compendia. BioData Mining. 2018;11(1):14-. doi: 10.1186/s13040-018-0175-7.

64. Tan J, Huyck M, Hu D, Zelaya RA, Hogan DA, Greene CS. ADAGE signature analysis: differential expression analysis with data-defined gene sets. BMC Bioinformatics. 2017;18(1):512-. doi: 10.1186/s12859-017-1905-4.

65. Recinos DA, Sekedat MD, Hernandez A, Cohen TS, Sakhtah H, Prince AS, et al. Redundant phenazine operons in *Pseudomonas aeruginosa* exhibit environment-dependent expression and differential roles in pathogenicity. Proceedings of the National Academy of Sciences. 2012;109(47):19420–5. doi: 10.1073/pnas.1213901109.

66. Mavrodi DV, Bonsall RF, Delaney SM, Soule MJ, Phillips G, Thomashow LS. Functional analysis of genes for biosynthesis of pyocyanin and phenazine-1-carboxamide from *Pseudomonas aeruginosa* PAO1. J Bacteriol. 2001;183(21):6454–65. doi: 10.1128/jb.183.21.6454-6465.2001.

67. Kanehisa M, Goto S. KEGG: kyoto encyclopedia of genes and genomes. Nucleic Acids Res. 2000;28(1):27–30. Epub 1999/12/11. doi: 10.1093/nar/28.1.27. PubMed PMID: 10592173; PubMed Central PMCID: PMCPMC102409.

68. Kanehisa M, Sato Y, Furumichi M, Morishima K, Tanabe M. New approach for understanding genome variations in KEGG. Nucleic Acids Res. 2019;47(D1):D590–d5. Epub 2018/10/16. doi: 10.1093/nar/gky962. PubMed PMID: 30321428; PubMed Central PMCID: PMCPMC6324070.

69. Kanehisa M. Toward understanding the origin and evolution of cellular organisms. Protein Sci. 2019;28(11):1947–51. Epub 2019/08/24. doi: 10.1002/pro.3715. PubMed PMID: 31441146; PubMed Central PMCID: PMCPMC6798127.

70. Grahl N, Demers EG, Lindsay AK, Harty CE, Willger SD, Piispanen AE, et al. Mitochondrial activity and Cyr1 are key regulators of Ras1 Activation of *C. albicans* virulence pathways. PLoS Path. 2015;11(8):e1005133. doi: 10.1371/journal.ppat.1005133.

71. Kwak MK, Ku M, Kang SO. Inducible NAD(H)-linked methylglyoxal oxidoreductase regulates cellular methylglyoxal and pyruvate through enhanced activities of alcohol dehydrogenase and methylglyoxal-oxidizing enzymes in glutathione-depleted *Candida albicans*. Biochimica et Biophysica Acta (BBA) - General Subjects. 2018;1862(1):18–39. Epub 2017/10/12. doi: 10.1016/j.bbagen.2017.10.003. PubMed PMID: 29017767.

72. Kwak MK, Ku M, Kang SO. NAD(+)-linked alcohol dehydrogenase 1 regulates methylglyoxal concentration in *Candida albicans*. FEBS Lett. 2014;588(7):1144–53. Epub 2014/03/13. doi: 10.1016/j.febslet.2014.02.042. PubMed PMID: 24607541.

73. Rampioni G, Falcone M, Heeb S, Frangipani E, Fletcher MP, Dubern JF, et al. Unravelling the genome-wide contributions of specific 2-Alkyl-4-quinolones and PqsE to quorum sensing in *Pseudomonas aeruginosa*. PLoS Pathog. 2016;12(11):e1006029. Epub 2016/11/17. doi: 10.1371/journal.ppat.1006029. PubMed PMID: 27851827; PubMed Central PMCID: PMCPMC5112799.

74. Lee J, Wu J, Deng Y, Wang J, Wang C, Wang J, et al. A cell-cell communication signal integrates quorum sensing and stress response. Nat Chem Biol. 2013;9(5):339–43. Epub 2013/04/02. doi: 10.1038/nchembio.1225. PubMed PMID: 23542643.

75. Meng X, Ahator SD, Zhang L-H. Molecular mechanisms of phosphate stress activation of *Pseudomonas aeruginosa* quorum sensing systems. mSphere. 2020;5(2):e00119–20. doi: 10.1128/mSphere.00119-20.

76. Schuster M, Lostroh CP, Ogi T, Greenberg EP. Identification, timing, and signal specificity of *Pseudomonas aeruginosa* quorum-controlled genes: a transcriptome analysis. J Bacteriol. 2003;185(7):2066–79. doi: 10.1128/jb.185.7.2066-2079.2003.

77. Déziel E, Gopalan S, Tampakaki AP, Lépine F, Padfield KE, Saucier M, et al. The contribution of MvfR to *Pseudomonas aeruginosa* pathogenesis and quorum sensing circuitry regulation: multiple quorum sensing-regulated genes are modulated without affecting *lasRI*, *rhlRI* or the production of N-acyl-l-homoserine lactones. Mol Microbiol. 2005;55(4):998–1014. doi: 10.1111/j.1365-2958.2004.04448.x.

78. Llamas MA, van der Sar A, Chu BCH, Sparrius M, Vogel HJ, Bitter W. A novel extracytoplasmic function (ECF) sigma factor regulates virulence in *Pseudomonas aeruginosa*. PLoS Path. 2009;5(9):e1000572. doi: 10.1371/journal.ppat.1000572.

79. Monds RD, Newell PD, Schwartzman JA, O’Toole GA. Conservation of the Pho regulon in *Pseudomonas fluorescens* Pf0-1. Appl Environ Microbiol. 2006;72(3):1910–24. doi: 10.1128/aem.72.3.1910-1924.2006.

80. Monds RD, Silby MW, Mahanty HK. Expression of the Pho regulon negatively regulates biofilm formation by *Pseudomonas aureofaciens* PA147-2. Mol Microbiol. 2001;42(2):415–26. doi: 10.1046/j.1365-2958.2001.02641.x.

81. Horwitz JP, Chua J, Noel M, Donatti JT, Freisler J. Substrates for cytochemical demonstration of enzyme activity. II. Some dihalo-3-indolyl phosphates and sulfates. Journal of Medical Chemistry. 1966;9(3):447. Epub 1966/05/01. doi: 10.1021/jm00321a059. PubMed PMID: 5960940.

82. Chamnongpol S, Groisman EA. Acetyl phosphate-dependent activation of a mutant PhoP response regulator that functions independently of its cognate sensor kinase. J Mol Biol. 2000;300(2):291–305. doi: https://doi.org/10.1006/jmbi.2000.3848.

83. Deretic V, Leveau JHJ, Mohr CD, Hibler NS. In vitro phosphorylation of AlgR, a regulator of mucoidy in *Pseudomonas aeruginosa*, by a histidine protein kinase and effects of small phospho-donor molecules. Mol Microbiol. 1992;6(19):2761–7. doi: 10.1111/j.1365-2958.1992.tb01455.x.

84. Hiratsu K, Nakata A, Shinagawa H, Makino K. Autophosphorylation and activation of transcriptional activator PhoB of *Escherichia coli* by acetyl phosphate in vitro. Gene. 1995;161(1):7–10. doi: https://doi.org/10.1016/0378-1119(95)00259-9.

85. Kim S-K, Wilmes-Riesenberg MR, Wanner BL. Involvement of the sensor kinase EnvZ in the in vivo activation of the response-regulator PhoB by acetyl phosphate. Mol Microbiol. 1996;22(1):135–47. doi: 10.1111/j.1365-2958.1996.tb02663.x.

86. Ikeh MAC, Kastora SL, Day AM, Herrero-de-Dios CM, Tarrant E, Waldron KJ, et al. Pho4 mediates phosphate acquisition in *Candida albicans* and is vital for stress resistance and metal homeostasis. Mol Biol Cell. 2016;27(17):2784–801. doi: 10.1091/mbc.E16-05-0266. PubMed PMID: 27385340.

87. Liu N-N, Flanagan PR, Zeng J, Jani NM, Cardenas ME, Moran GP, et al. Phosphate is the third nutrient monitored by TOR in *Candida albicans* and provides a target for fungal-specific indirect TOR inhibition. Proceedings of the National Academy of Sciences. 2017;114(24):6346–51. doi: 10.1073/pnas.1617799114.

88. Lev S, Djordjevic JT. Why is a functional PHO pathway required by fungal pathogens to disseminate within a phosphate-rich host: A paradox explained by alkaline pH-simulated nutrient deprivation and expanded PHO pathway function. PLoS Path. 2018;14(6):e1007021-e. doi: 10.1371/journal.ppat.1007021. PubMed PMID: 29928051.

89. Liu N-N, Uppuluri P, Broggi A, Besold A, Ryman K, Kambara H, et al. Intersection of phosphate transport, oxidative stress and TOR signalling in *Candida albicans* virulence. PLoS Path. 2018;14(7):e1007076. doi: 10.1371/journal.ppat.1007076.

90. Urrialde V, Prieto D, Pla J, Alonso-Monge R. The *Candida albicans* Pho4 transcription factor mediates susceptibility to stress and influences fitness in a mouse commensalism model. Front Microbiol. 2016;7(1062). doi: 10.3389/fmicb.2016.01062.

91. Crocker AW, Harty CE, Hammond JH, Willger SD, Salazar P, Botelho NJ, et al. *Pseudomonas aeruginosa* ethanol oxidation by AdhA in low oxygen environments. J Bacteriol. 2019:JB.00393-19. doi: 10.1128/jb.00393-19.

92. Mern DS, Ha S-W, Khodaverdi V, Gliese N, Görisch H. A complex regulatory network controls aerobic ethanol oxidation in *Pseudomonas aeruginosa*: indication of four levels of sensor kinases and response regulators. Microbiology. 2010;156(5):1505–16. doi: doi:10.1099/mic.0.032847-0.

93. Hornby JM, Jensen EC, Lisec AD, Tasto JJ, Jahnke B, Shoemaker R, et al. Quorum sensing in the dimorphic fungus *Candida albicans* is mediated by farnesol. Appl Environ Microbiol. 2001;67(7):2982–92. PubMed PMID: 11425711.

94. Qi Y, Kobayashi Y, Hulett FM. The *pst* operon of *Bacillus subtilis* has a phosphate-regulated promoter and is involved in phosphate transport but not in regulation of the pho regulon. J Bacteriol. 1997;179(8):2534–9. doi: 10.1128/jb.179.8.2534-2539.1997.

95. Nikata T, Sakai Y, Shibata K, Kato J, Kuroda A, Ohtake H. Molecular analysis of the phosphate-specific transport (*pst*) operon of *Pseudomonas aeruginosa*. MGG Molecular & General Genetics. 1996;250(6):692–8. doi: 10.1007/BF02172980.

96. Madhusudhan KT, McLaughlin R, Komori N, Matsumoto H. Identification of a major protein upon phosphate starvation of *Pseudomonas aeruginosa* PAO1. J Basic Microbiol. 2003;43(1):36–46. doi: 10.1002/jobm.200390002.

97. Kim H-Y, Schlictman D, Shankar S, Xie Z, Chakrabarty AM, Kornberg A. Alginate, inorganic polyphosphate, GTP and ppGpp synthesis co-regulated in *Pseudomonas aeruginosa*: implications for stationary phase survival and synthesis of RNA/DNA precursors. Mol Microbiol. 1998;27(4):717–25. doi: 10.1046/j.1365-2958.1998.00702.x.

98. Gallarato LA, Sanchez DG, Olvera L, Primo ED, Garrido MN, Beassoni PR, et al. Exopolyphosphatase of *Pseudomonas aeruginosa* is essential for the production of virulence factors, and its expression is controlled by NtrC and PhoB acting at two interspaced promoters. Microbiology. 2014;160(Pt_2):406-17. doi: 10.1099/mic.0.074773-0.

99. . Almeida LGd, Ortiz JH, Schneider RP, Spira B. *phoU* Inactivation in *Pseudomonas aeruginosa* enhances accumulation of ppGpp and polyphosphate. Appl Environ Microbiol. 2015;81(9):3006–15. doi: 10.1128/AEM.04168-14.

100. Bertani G. Studies on lysogenesis. I. The mode of phage liberation by lysogenic Escherichia coli. J Bacteriol. 1951;62(3):293–300. Epub 1951/09/01. PubMed PMID: 14888646; PubMed Central PMCID: PMC386127.

101. Shanks RM, Caiazza NC, Hinsa SM, Toutain CM, O’Toole GA. *Saccharomyces cerevisiae*-based molecular tool kit for manipulation of genes from gram-negative bacteria. Appl Environ Microbiol. 2006;72(7):5027–36. PubMed PMID: 16820502.

102. Gibson DG, Glass JI, Lartigue C, Noskov VN, Chuang R-Y, Algire MA, et al. Creation of a bacterial cell controlled by a chemically synthesized genome. Science. 2010;329(5987):52-6. doi: 10.1126/science.1190719.

103. Gibson DG, Young L, Chuang R-Y, Venter JC, Hutchison CA, Smith HO. Enzymatic assembly of DNA molecules up to several hundred kilobases. Nat Methods. 2009;6(5):343–5. doi: 10.1038/nmeth.1318.

104. Team RDC. R: A language and environemnt for statistical computing. Vienna, Austria: R Foundation for Statistical Computing; 2010.

105. Wickham H. ggplot2: Elegent Graphics for Data Analysis. New York: Springer-Verlag; 2016.

106. Robinson M, McCarthy D, Smyth G. edgeR: a Bioconductor package for differential expression analysis of digital gene expression data. Bioinformatics. 2010;26(1):139–40.

107. Tenenbaum D. KEGGREST: Client-side REST access to KEGG. 2018.

108. Zuguang G, Eils R, Schlesner M. Complex heatmaps reveal patterns and correlations in multidimensional genomic data. Bioinformatics. 2016.

109. Miller JH. A Short Course in Bacterial Genetics: Cold Spring Harbor Press; 1992. 456 p.

110. Sacks LE. A pH gradient agar plate. Nature. 1956;178(4527):269-70. doi: 10.1038/178269a0.

## References

1. Rahme LG, Stevens EJ, Wolfort SF, Shao J, Tompkins RG, Ausubel FM. Common virulence factors for bacterial pathogenicity in plants and animals. Science. 1995;268(5219):1899–902. PubMed PMID: 7604262.

2. Tan J, Doing G, Lewis KA, Price CE, Chen KM, Cady KC, et al. Unsupervised Extraction of Stable Expression Signatures from Public Compendia with an Ensemble of Neural Networks. Cell Syst. 2017;5(1):63–71 e6. doi: 10.1016/j.cels.2017.06.003. PubMed PMID: 28711280; PubMed Central PMCID: PMCPMC5532071.

3. Liberati NT, Urbach JM, Miyata S, Lee DG, Drenkard E, Wu G, et al. An ordered, nonredundant library of Pseudomonas aeruginosa strain PA14 transposon insertion mutants. Proc Natl Acad Sci U S A. 2006;103(8):2833–8. Epub 2006/02/16. doi: 10.1073/pnas.0511100103. PubMed PMID: 16477005; PubMed Central PMCID: PMCPMC1413827.

4. Dietrich LEP, Price-Whelan A, Petersen A, Whiteley M, Newman DK. The phenazine pyocyanin is a terminal signalling factor in the quorum sensing network of Pseudomonas aeruginosa. Mol Microbiol. 2006;61(5):1308–21. doi: 10.1111/j.1365-2958.2006.05306.x.

5. Sakhtah H, Koyama L, Zhang Y, Morales DK, Fields BL, Price-Whelan A, et al. The *Pseudomonas aeruginosa* efflux pump MexGHI-OpmD transports a natural phenazine that controls gene expression and biofilm development. Proc Natl Acad Sci U S A. 2016;113(25):E3538–47. doi: 10.1073/pnas.1600424113.

6. Crocker AW, Harty CE, Hammond JH, Willger SD, Salazar P, Botelho NJ, et al. *Pseudomonas aeruginosa* ethanol oxidation by AdhA in low oxygen environments. J Bacteriol. 2019:JB.00393-19. doi: 10.1128/jb.00393-19.

7. Fonzi WA, Irwin MY. Isogenic strain construction and gene mapping in Candida albicans. Genetics. 1993;134(3):717–28. Epub 1993/07/01. PubMed PMID: 8349105; PubMed Central PMCID: PMCPMC1205510.

8. Chen AI, Dolben EF, Okegbe C, Harty CE, Golub Y, Thao S, et al. *Candida albicans* ethanol stimulates *Pseudomonas aeruginosa* WspR-controlled biofilm formation as part of a cyclic relationship involving phenazines. PLoS Path. 2014;10(10):e1004480- e. doi: 10.1371/journal.ppat.1004480.

9. Glasser NR, Kern SE, Newman DK. Phenazine redox cycling enhances anaerobic survival in Pseudomonas aeruginosa by facilitating generation of ATP and a proton-motive force. Mol Microbiol. 2014;92(2):399–412. Epub 2014/03/19. doi: 10.1111/mmi.12566. PubMed PMID: 24612454.

10. Feinbaum RL, Urbach JM, Liberati NT, Djonovic S, Adonizio A, Carvunis AR, et al. Genome-wide identification of *Pseudomonas aeruginosa* virulence-related genes using a *Caenorhabditis elegans* infection model. PLoS Pathog. 2012;8(7):e1002813. doi: 10.1371/journal.ppat.1002813. PubMed PMID: 22911607; PubMed Central PMCID: PMCPMC3406104.

11. Quesada JM, Otero-Asman JR, Bastiaansen KC, Civantos C, Llamas MA. The activity of the *Pseudomonas aeruginosa* virulence regulator σVreI is modulated by the anti-σ factor VreR and the transcription factor PhoB. Front Microbiol. 2016;7:1159-. doi: 10.3389/fmicb.2016.01159.

12. Shanks RM, Caiazza NC, Hinsa SM, Toutain CM, O’Toole GA. *Saccharomyces cerevisiae*-based molecular tool kit for manipulation of genes from gram-negative bacteria. Appl Environ Microbiol. 2006;72(7):5027–36. PubMed PMID: 16820502.

13. Qiu D, Damron FH, Mima T, Schweizer HP, Yu HD. PBAD-based shuttle vectors for functional analysis of toxic and highly regulated genes in Pseudomonas and Burkholderia spp. and other bacteria. Appl Environ Microbiol. 2008;74(23):7422–6. Epub 2008/10/14. doi: 10.1128/aem.01369-08. PubMed PMID: 18849445; PubMed Central PMCID: PMCPMC2592904.

14. Damron FH, Qiu D, Yu HD. The *Pseudomonas aeruginosa* sensor kinase KinB negatively controls alginate production through AlgW-dependent MucA proteolysis. J Bacteriol. 2009;191(7):2285–95. Epub 2009/01/23. doi: 10.1128/JB.01490-08. PubMed PMID: 19168621.

15. Hoang TT, Karkhoff-Schweizer RR, Kutchma AJ, Schweizer HP. A broad-host-range Flp-FRT recombination system for site-specific excision of chromosomally-located DNA sequences: application for isolation of unmarked *Pseudomonas aeruginosa* mutants. Gene. 1998;212(1):77–86. PubMed PMID: 9661666.

